# Lifetime fitness and annual survival are heritable and highly genetically correlated in a wild primate population

**DOI:** 10.1101/2025.11.13.688343

**Authors:** Beniamino Tuliozi, Elizabeth A. Archie, Jenny Tung, Susan Alberts

**Author notes:** Address: Duke University, Department of Biology, Box 90383, Durham, NC 27708, USA.

## Abstract

The additive genetic variance (V_A_) of fitness quantifies the expected response to selection; lifetime breeding success (LBS) is an effective metric of fitness in animal populations. However, data on LBS are relatively rare for wild populations of long-lived species, while time-limited metrics such as annual survival or fertility are more readily available. The magnitude of V_A_ for these time-limited metrics, and the degree to which they are genetically correlated with LBS, remains unclear in most cases. Here we estimated the V_A_ and heritability (h^2^) of LBS and four time-limited fitness metrics in wild female baboons in Kenya. The most highly heritable metrics were LBS (h^2^=0.25) and annual survival (h^2^=0.23). Notably, all the V_A_ for LBS was attributable to survival to first successful reproduction. Furthermore, all fitness metrics examined were highly genetically correlated with each other, supporting the potential use of time-limited metrics where LBS data are limited. Our analyses predicted faster phenotypic evolution than we observed, raising the possibility that environmental effects have masked responses to selection (“cryptic evolution”) or that social effects inflate estimated V_A_. Together, our findings reveal a substantial genetic contribution to variation in survival, and in turn, to fitness and contemporary evolution in a long-lived animal.

## INTRODUCTION

Individual fitness is the complex outcome of multiple influences, external and internal to the organism. Environmental, social, and maternal effects, as well as random chance, can all greatly influence fitness (Schroeder *et al*. 2012; Sæther & Engen 2015; Snyder & Ellner 2018; Snyder-Mackler *et al*. 2020). At the same time, fitness is also the emergent outcome of individual traits that contribute to fertility and survival, and that often have a genetic basis (Wolak *et al*. 2018; Warrington *et al*. 2024). The heritable component of these traits constitutes the genetic basis of fitness, which in turn determines if, and at what rate, evolutionary change can occur in natural populations (Fisher 1930; Price 1970; Price 1972; Hendry *et al*. 2018; Bonnet *et al*. 2019a). Quantifying the genetic basis of fitness is essential for assessing a population’s potential for adaptive evolution in the face of natural selection, and for understanding how evolution has shaped a species’ life history and behaviour.

In recent years, interest in the additive genetic variance (V_A_) of fitness has soared, thanks to the emerging availability of datasets from long-term studies that enable the V_A_ of fitness to be estimated in nature (Sheldon *et al*. 2022). At the same time, the quantitative genetic techniques used in the analysis of such datasets have continued to be refined (Postma 2014; de Villemereuil *et al*. 2016; de Villemereuil *et al*. 2018; Bonnet *et al*. 2019a). Because individual fitness is an emergent property of many traits, it is typically treated as highly polygenic. This representation makes variation in fitness amenable to analysis with mixed effect models that partition trait variance by attributing variance to genetic effects via an additive genetic relationship matrix (‘animal model’; Kruuk (2004); Wilson *et al*. (2010)). A key advantage of these models is that they can account for the effect of non-genetic as well as genetic predictors.

Lifetime breeding success (LBS, sensu Bonnet *et al*. (2022)), or the number of progeny produced over the course of an individual lifetime, is a common metric of individual lifetime fitness (Hendry *et al*. 2018; Young *et al*. 2023). However, for a precise and unbiased measure of LBS, the entire life history of most individuals in all considered cohorts should be recorded – data that are difficult to obtain, especially for long-lived species in the wild. Furthermore, both the physical and the social environment have the potential to affect fitness metrics, but their influence is difficult to capture accurately when LBS reflects the composite contribution of many years. Thus, LBS is not an appropriate metric for investigating selective pressures that vary on time-scales shorter than an adult’s lifespan because LBS can be too coarse to capture fine-scale variation in the selective environment (Coulson *et al*. 2006; Scranton *et al*. 2016).

For these reasons, studies on natural populations of long-lived animals have often employed alternatives to LBS. One set of alternatives involves using proxy traits thought to be correlated with LBS, such as age at first reproduction, birth weight, adult body size, or foraging rate (Hendry *et al*. 2018; Alif *et al*. 2022). However, such proxy traits are indirect measures of fitness that are usually highly system-specific and thus difficult to generalize. Moreover, the strength of their actual correlation with fitness is often untested (Hendry *et al*. 2018). A second set of alternatives involves using time-limited fitness metrics measured over specific intervals of the individual lifetime, such as annual survival or annual fertility (Qvarnström *et al*. 2006; Postma 2014). These metrics directly measure fitness components while also capturing influences on fitness and selection that operate on timescales shorter than the full life course, which may include variation in the social environment, habitat quality, and annual environmental variation (Viblanc *et al*. 2022). Hence, using these metrics has the potential advantage to improve our understanding of the genetic components of fitness in the wild (Wilson 2008; de Villemereuil *et al*. 2018).

### The genetic correlation between lifetime and time-limited fitness metrics

To provide useful insight into the genetic basis of fitness, time-limited fitness metrics must be genetically correlated with lifetime fitness. Genetic correlations between different life history traits have been investigated in several taxa, especially invertebrates (see the meta-analysis by Chang *et al*. (2024)), usually with the purpose of determining the presence of trade-offs or life-history constraints (Morrissey *et al*. 2012; Teplitsky *et al*. 2014; Chang *et al*. 2024). However, few studies have estimated the genetic correlations between time-limited and lifetime fitness metrics (Schroeder *et al*. 2012; Walling *et al*. 2014; Wolak *et al*. 2018). Part of the reason is the historic lack of appropriate methods for dealing with non-Gaussian distributed traits (Morrissey 2015; de Villemereuil *et al*. 2016). Recent methodological advances, however, now enable V_A_ estimation for a more general set of traits, including those that follow zero-inflated Poisson distributions (e.g. many measures of LBS) or binomial distributions (e.g. annual survival) (Bonnet *et al*. 2019a; Morrissey & Bonnet 2019; Bonnet *et al*. 2022). While several recent studies have compared different fitness metrics with the specific purpose of determining their interchangeability, their respective biological relevance, and their relative merits (Dobson *et al*. 2020; Alif *et al*. 2022; Van de Walle *et al*. 2022; Viblanc *et al*. 2022; Young *et al*. 2023), to date no comprehensive estimates of genetic correlations between lifetime fitness and time-limited fitness metrics have been conducted. Estimating these genetic correlations is important because they reveal whether different fitness metrics behave similarly when used to quantify selection and evolutionary change. Empirically testing the interchangeability of time-limited and lifetime fitness metrics by estimating their genetic correlations will thus provide an evidence-based foundation for interpreting results obtained with time-limited fitness metrics. Such a foundation is especially important for wild study systems where recording LBS is not always possible.

### Study aims

We estimated additive genetic variance (V_A_) and genetic covariance for lifetime and time-limited fitness metrics in the wild baboons of the Amboseli basin of southern Kenya. The Amboseli baboons are ideally suited to compare time-limited and lifetime fitness metrics because multiple generations of individuals have been followed continuously for five decades, producing numerous complete cohorts for unbiased estimates of female lifetime breeding success. Here, we focus on females because male dispersal makes estimates of male LBS considerably more difficult to obtain. In addition, the wealth of detailed complementary information available on this population gives us the rare opportunity to estimate the effects of time-varying social variables (group identity, group size) and environmental conditions (rainfall, habitat quality) on time-limited fitness metrics. Notably, in a comparison of multiple mammal and bird studies, the Amboseli baboons were found to have one of the highest estimates of additive genetic variance in relative lifetime fitness (V_A_(*w*), (Bonnet *et al*. (2022)). This result makes them a good candidate for further investigation of V_A_ across both lifetime and time-limited fitness metrics.

Our first aim was to estimate the magnitude of V_A_ for both time-limited and lifetime fitness metrics for female baboons in this population. To achieve this aim, we fitted univariate animal models for both LBS and four time-limited measures of fitness: annual survival, annual breeding success, annual fitness, and triannual breeding success. Our second aim was to calculate genetic correlations between these fitness metrics, with the ultimate objective of determining the interchangeability of time-limited and lifetime fitness metrics in Amboseli baboons. To achieve this aim, we fitted bivariate animal models for all pairwise combinations of the above-mentioned fitness metric. These models allowed us to estimate the genetic covariance(s) between each pair of traits (i.e. the degree to which different fitness components have a shared genetic basis), and to identify the largest source(s) of genetic variance in fitness in this population. A positive genetic correlation between two fitness metrics indicates that a given genotype produces similarly high (or low) values for both metrics, while a negative correlation indicates antagonistic pleiotropy (a genetic trade-off), where high values for one metric predict low values for the other. While some genetic trade-offs have been well-documented, meta-analytic findings show weak to mixed evidence for antagonistic trade-offs between reproduction and survival across a wide range of animal taxa (Chang *et al*. (2024); Winder *et al*. (2025), but see Ryan *et al*. (2024)). Thus, we predicted either zero or positive genetic correlations between the measures we considered (Walling *et al*. 2014; Dobson *et al*. 2020). Our approach therefore allowed us to evaluate the value and feasibility of applying time-limited fitness metrics in evolutionary studies, their ability to evolve independently, and their relative importance for selection and evolutionary change in this population.

## METHODS

### Study population and subjects

The baboon population in the Amboseli basin of southern Kenya has been the subject of ongoing longitudinal data collection since 1971 (Alberts & Altmann 2012). The population consists of primarily yellow baboons (*Papio cynocephalus*) that have experienced historical and recent admixture with anubis baboons (*P. anubis*) (Wall *et al*. 2016; Vilgalys *et al*. 2022). The Amboseli ecosystem is a highly seasonal semi-arid savanna: the ‘hydrological year’ in Amboseli (the ‘hydroyear’) starts on the 1^st^ of November each year with the onset of the rainy season and ends on October 31^st^ of the following calendar year. Females conceive and give birth throughout the year, with only mild seasonal variation (Gesquiere *et al*. 2024).

Baboons in Amboseli live in stable social groups, ranging from 15 to 130 individuals. A subset of the social groups in Amboseli, hereafter ‘study groups’, are each observed several times a week. Data are collected on demographic and life history events (births, deaths, maturations, dispersals) and social behavior. Because female baboons are philopatric, females in study groups are followed from birth until death, unless they were alive before the study groups were established (left-truncated) or belonged to a group where monitoring stopped (right-censored). Our analyses included all life history events that occurred for all subjects when they were in a study group. The pedigree was based on observations of pregnancies and births, augmented by genetic analysis (Galezo *et al*. 2022; McLean *et al*. 2023). The full pedigree consists of 1918 individuals, including 1789 individuals with known mothers and 651 individuals with known fathers. The pedigree was trimmed (individuals unrelated to the study subjects were eliminated from further analysis) using the *prunePed* function of the nadiv package (Wolak 2012), resulting in a final pedigree for annual fitness metrics that included 932 individuals: 828 with known mothers, 354 with known fathers, 47 full sibling pairs, 1271 maternal half-sibling pairs, and 658 paternal half-sibling pairs (Table 1). The research in this study was approved by the Institutional Animal Care and Use Committee (IACUC) at Duke University (current protocol no. A003-24-01) and adhered to the laws and guidelines of the Kenyan government.

**Table 1.** Fitness metrics used in our study: definitions, sample sizes, data distributions, and pedigree.

|  | **Fitness metric** | **Definition** | **Sample size** | **Distribution** | **Trimmed pedigree** |
| --- | --- | --- | --- | --- | --- |
| Time-limited metrics | Annual breeding success | Number of offspring born during each hydroyear of a female’s life that survived to at least 365 days; binary (0 or 1 for each female-hydroyear) | 806 females, 5896 female-hydroyears | Binomial distribution analysed with threshold GLMM | • 932 individuals<br>• 828 known mothers, 354 known fathers<br>• 47 full sib pairs<br>• 658 paternal half sib pairs<br>• 1271 maternal half sib pairs |
|  | Annual survival | Binary variable, with 1 for every female-hydroyear when the individual was alive, and 0 for the female-hydroyear of its death | 806 females, 5896 female-hydroyears | Binomial distribution analysed with threshold GLMM | Same as for annual breeding success |
|  | Annual fitness | Sum of annual breeding success and annual survival (ordinal values of 0, 1, or 2 for each female-hydroyear) | 806 females, 5896 female-hydroyears | Ordinal distribution analysed with ordinal GLMM | Same as for annual breeding success |
|  | Triannual breeding success | Number of offspring born during each three-year interval (triennium) of a female’s life that survived to at least 365 days; ordinal values of 0, 1, 2, 3 for each female-triennium | 790 females<sup>a</sup>, 2036 female-triennia | Ordinal distribution analysed with ordinal GLMM | • 918 individuals,<br>• 817 known mothers, 354 known fathers,<br>• 47 full sib pairs,<br>• 658 paternal half sib pairs,<br>• 1268 maternal half sib pairs |
| Lifetime metrics | LBS | Total number of offspring produced during the lifetime of a female, counting only infants that survived at least 365 days | 265 females | Zero-inflated Poisson distribution analyzed with zero-inflated Poisson GLMM | • 345 individuals,<br>• 293 known mothers, 107 known fathers,<br>• 3 full sib pairs,<br>• 131 paternal half sib pairs,<br>• 363 maternal half sib pairs |
|  | Binary LBS | Whether or not the female ever produced an offspring that survived to 365 days | 265 females | Binomial distribution analyzed with threshold GLMM | Same as for total LBS |

<sup>a</sup> The triennial dataset is smaller than the annual one because more individuals are censored as a result of the fact that the triennium is longer than the hydroyear: it is thus more common for a triennium to be censored at the endpoint of this study.

### Quantitative genetic analyses of fitness metrics

#### Response variables and datasets

To estimate the genetic variance associated with fitness metrics we used a variance-partitioning approach called the animal model (Wilson *et al*. 2010). The response variables in the animal model are vectors of trait values (in this case, fitness metrics). The model estimates the fraction of trait variance (phenotypic variance) that can be explained by pedigree-based genetic relationships between individuals (Kruuk 2004). We fitted six univariate animal models, each with one of the following response variables: *i*) *annual breeding success, ii) annual survival, iii) annual fitness, iv) triannual breeding success* (i.e., breeding success across three consecutive years)*, v) lifetime breeding success* (*LBS*) and *vi*) *binary lifetime breeding success* (*binary LBS*; see below and Table 1 for definitions).

For all fitness metrics that included the number of offspring produced by a female (i.e., all but annual survival), we counted only infants that survived at least 365 days (hereafter “surviving offspring”). This age corresponds closely to near-complete weaning and the earliest age at which offspring can survive the death of their mother in this population (Altmann 1998; Alberts & Altmann 2003; Altmann & Alberts 2003). Using this age as our metric of offspring production is analogous to using counts of offspring that survive to various post-zygotic stages in other species (e.g., Moiron *et al*. (2022); Viblanc *et al*. (2022); Dunning *et al*. (2023)). While zygote-to-zygote measures are theoretically preferred (Arnold & Wade 1984), they are not commonly used because of practical issues in recording zygote-to-zygote life histories (Hendry *et al*. 2018; Wolak *et al*. 2018). Moreover, lifetime fitness metrics are better predictors of individual genetic contributions to the future population when measured at later stages (Alif *et al*. 2022). For all time-limited fitness metrics, we excluded female-hydroyears (or female triennia) in which a hydroyear included less than 90 days of data, unless it represented the hydroyear in which the female was born or died.

##### Annual breeding success

For annual breeding success, our dataset included 806 female baboons that lived in wild-feeding (i.e., non-provisioned) study groups between 1971 and 2022, resulting in a total dataset of 5896 female-hydroyears (see Table 1 for dataset descriptions). We modeled annual breeding success as the number of surviving offspring born during each hydroyear of a female’s life. No females in the dataset produced two surviving offspring during the same hydroyear. Consequently, annual breeding success defaults to a binary trait, corresponding to a value of 1 for female-hydroyears in which an offspring that survived to 365 days was produced, and 0 for female-hydroyears in which no surviving offspring was produced. Note that we measured a female’s annual breeding success in all years of her life, including years in which she was immature; females that died before adulthood therefore only had annual breeding success values of 0. This approach ensured that we produced an unbiased fitness metric that includes a crucial component of the true variance in annual breeding success, as approximately 48% of females in this study population do not reach reproductive age.

##### Annual survival

For annual survival, we used the same dataset as for annual breeding success (N=806 females during 5,896 female-hydroyears; Table 1). We modeled annual survival as a binary variable: 1 for every full hydroyear that a female survived, including her birth hydroyear, and 0 for the hydroyear of death.

##### Annual fitness

We modeled annual fitness as a sum of annual breeding success and annual survival (Brommer et al. 2007a; Ranke et al. 2017). Thus, each female baboon was assigned one of three ranked values during each hydroyear of her life: 0 (hydroyear of death), 1 (alive for the full hydroyear), or 2 (alive for the full hydroyear and produced an offspring).

##### Triannual breeding success

We included a triannual metric of breeding success (fertility) because an annual period is short relative to the breeding cycle of female baboons (Gesquiere *et al*. 2018) and therefore might not adequately capture differences in female fertility. For triannual breeding success, our dataset included 790 females during 2036 female-triennia (Table 1). This dataset is somewhat smaller than the dataset for the annual metrics because incomplete intervals occur more frequently for three-year than one-year intervals. We modeled triannual breeding success as the number of surviving offspring that a female produced within each three-year interval. Each female baboon was assigned one of four ordinal values during each triennium of her life: 0, 1, 2 or 3 depending upon how many surviving offspring she produced in that triennium. The first three-year interval began at birth, the second at 3 years of age, the third at 6 years of age, and so on; we included triennia in which the female died. While the annual metrics refer to hydroyears and thus have the same timeframe for all individuals, triannual breeding success was individually calculated, starting on the day of each female’s birth; the last interval ended on the individual’s death.

##### Lifetime breeding success (LBS)

We modeled LBS as the total number of surviving offspring produced during the lifetime of an individual (i.e., offspring that survived at least 365 days). For LBS, our dataset comprised 265 female baboons (Table 1). This dataset is smaller than the datasets for annual and triannual metrics because, to provide accurate, unbiased, and comparable measures of LBS, we required that both each subject’s life history records and those of her age cohort be complete (but see Moiron *et al*. (2022) for an alternative). We therefore included only female baboons i) who were followed from birth to death, ii) for whom the birth dates and survival status of all her offspring were known, and iii) who were born before hydroyear 2002, because all but one female born prior to 2002 had died by the time of this analysis. We also excluded all baboons born before systematic observations began in 1971, and all individuals that were not followed throughout their entire life (e.g., because their study groups were dropped).

##### Binary lifetime breeding success (binary LBS)

Our preliminary analysis revealed that most of the genetic variance in LBS was linked to whether a female produced at least one infant during her life (which in turn was almost entirely predicted by whether she survived to adulthood; see “Univariate models” below). Therefore, we modeled lifetime breeding success a second time as a binary outcome (binary LBS), using the same dataset as the LBS analysis (Table 1). We assigned 1 to all females who produced at least one surviving infant, and 0 to all females that did not.

#### Fixed and random effects

We included several biologically relevant non-genetic random and fixed effects in our models based on previous evidence that they influence fitness-related outcomes (McLean *et al*. 2019; Campos *et al*. 2020). We z-score transformed all continuous covariates to make comparisons between estimates more intuitive (see Supplementary Methods S1). Because missing pedigree links and admixture-related variation in genetic ancestry could also affect our estimates of V_A_ but are documented only for a subset of genetically well-characterized individuals, we also explored their potential impact in similar models with smaller data sets (see Supplementary Methods S2-S3, Supplementary Table S1).

##### Annual breeding success, annual fitness, *and* annual survival

The three annual fitness metrics were modeled using the following fixed effects (see Supplementary Methods S1 for details on each fixed effect): i) the subject’s age, based on her birthday in that hydroyear, and ii) age squared; iii) average social group size (count of adult females) during that hydroyear and iv) average group size squared; v) habitat quality during the hydroyear, and vi) rainfall anomaly for the previous hydroyear. We also included random effects of: i) the permanent environment, representing individual non-genetic repeatability; ii) maternal identity (some values were missing as the mothers of 32 females were unknown); iii) hydroyear, to account for random differences in environmental conditions between hydroyears; and iv) hydroyear-group, to capture variance associated with random variation between the environments of different social groups.

##### Triannual breeding success

Triannual breeding success was modeled with fixed effects averaged over three-year intervals (triennia) in the life of each female baboon (see Supplementary Methods S1): i) triennium, as a measure of age (e.g., triennium 1 covers birth to 3 years of age; triennium 2 corresponds to 3 – 6 years of age, and so on); ii) triennium squared; iii) group size; iv) group size squared; v) habitat quality where the majority of the triennium was spent; and vi) rainfall anomaly over a 3-year period starting 365 days before the start of the triennium. Additional random effects included i) permanent environment (individual non-genetic repeatability) and ii) maternal identity. Because each sequence of triennia was calculated uniquely for every individual, it was not possible to fit random effects associated with specific time units, like hydroyear or group-hydroyear.

##### LBS and Binary LBS

Many dramatic changes can occur in the physical and social environment of a female baboon during her long life; averaging these effects over the entire lifespan is potentially meaningless. Therefore, we limited the fixed effects for LBS to environmental influences that could be compared across individuals of any age. We did not include measures of early life adversity associated with individual characteristics (e.g., dominance rank, social connectedness) because such characteristics can also be influenced by genotype, which could therefore bias our fitness estimates (Wilson 2008). Our fixed effects therefore included: i) rainfall anomaly a year before birth; ii) habitat quality at birth; and iii) hydroyear of birth as a linear covariate. Additional random effects included i) maternal identity and ii) hydroyear of birth. Following Bonnet *et al*. (2022), hydroyear of birth was fitted both as a fixed and a random effect, to account for both linear and random time-linked variation during the study period.

#### Implementation of animal models

We ran all our mixed-effects animal models in a Bayesian framework using the MCMCglmm package (Hadfield 2010) within the R environment (R Core Team 2013). We tested convergence by visually assessing the posterior density distributions and chains, and by using the Heidelberger and Welch convergence test implemented in the coda package in R (Plummer *et al*. 2007). To obtain an appropriate effective sample size for characterizing the posterior distribution we ran our models with different combinations of thinning, number of iterations, and burn in (reported in Supplementary Table S2). In models involving LBS and bivariate models involving binary LBS we detected under/overflow and constrained the latent values accordingly (models with non-truncated latent variables returned almost identical results) (Hadfield 2010; Hadfield *et al*. 2013; Regan *et al*. 2019).

Since all of our fitness metrics produced non-Gaussian data distributions, we reported our results on the liability scale (a representation of the phenotype as a normally distributed “latent trait;” the value of this latent trait affects the probability of a positive outcome for an observed binary trait, or higher count values for ordinal traits) (de Villemereuil *et al*. 2016; de Villemereuil 2018). We also back-transformed them to the observed scale (i.e., the scale of the original data) using the *QGparams* and *QGmvparams* functions in QGglmm package (de Villemereuil *et al*. 2016). Back-transformation allows us to draw evolutionary inferences and to include fixed effects in the estimate of genetic parameters, as it considers the true distribution of the trait (de Villemereuil *et al*. 2018; Bonnet *et al*. 2019a). For more information on data distribution scales see Supplementary Methods S4 (“Glossary”).

We provide the 95% credible intervals (CI) built on model iterations after burn-in for both fixed and random effects. Fixed effects were highlighted if the 95% CI did not include zero. We considered variances attributed to random effects to be non-zero if the lower end of their 95% CI exceeded 0.001 (Bonnet *et al*. 2022; Dobson *et al*. 2023). We highlighted genetic correlations for which the 95% CI for the correlation coefficient did not overlap zero.

#### Univariate models

Our first aim was to fit univariate animal models to estimate the additive genetic variance (V_A_) of each fitness metric alone. We implemented models for (i) binomially distributed fitness metrics, (annual breeding success, annual survival, binary LBS) using the ‘threshold’ family specification in MCMCglmm; (ii) ordinal fitness metrics (annual fitness, triannual breeding success) using MCMCglmm’s ‘ordinal’ family specification; and (iii) our single zero-inflated Poisson distributed metric (LBS), using the ‘zipoisson’ family specification. We used parameter expanded priors for all models (de Villemereuil *et al*. 2013) and ran them to obtain minimum effective sample sizes from the posterior distribution of >9000 for all fixed effects and >6800 for all random effects, with autocorrelation below 0.1 for all variances. To back-transform variance estimates to the observed scale we used the ‘binom1.probit’ model specification from *QGparams* for binomially distributed variables (results did not change when using the “logit” link) and the ‘ordinal’ specification for ordinal metrics. For ordinal variables, however, we discuss the liability scale results because they produce a single estimate of genetic variance, rather than the variance-covariance matrix generated by back-transformed values, and are generally the preferred ones for interpretation (de Villemereuil 2018). For LBS, we first back-transformed estimates of variance components separately for the zero-inflated and the Poisson components, which is useful since they have distinct biological meaning (Moiron *et al*. 2022). The zero-inflated component represents whether or not the female ever produced an offspring, which is strongly shaped by juvenile survival, while the Poisson component represents the number of offspring produced (i.e. adult reproductive success). We also obtained a combined, meaningful estimate of the total observed genetic variance and total heritability for the LBS trait (including both components together) using the ‘ZIPoisson.log.logit’ specification for the *QGmvparams* function (kindly provided by P. de Villemereuil) within the “compound” branch of the QGglmm package.

### Bivariate models: response variables, effects structure, and implementation

To estimate genetic correlations between our fitness metrics, we fitted 15 bivariate animal models, featuring all possible pairs of fitness metrics as bivariate response variables. We used the same datasets, response variables, and suites of fixed effects for the bivariate analyses as for the univariate analyses. In the bivariate analyses, however, the random effect structures included both the random effects and, when appropriate, their covariances (see Supplementary Methods S5-S6). All bivariate models were implemented in MCMCglmm using the families of the univariate models that composed them. Back-transformation for bivariate models was performed in the same way as for univariate models but with the *QGmvparams* function from QGglmm package; we used 120-500 iterations per bivariate model (Biquet *et al*. 2022). For models of the “ordinal” family no specification is currently available for *QGmvparams* so we used “binom1.probit”. We iterated all bivariate models enough times to obtain minimum effective sample sizes of 1114 for random effects and 1691 for fixed effects (see Supplementary Methods S6, Table S2). To further test the robustness of our parameter estimates we ran a subset of bivariate models with a permuted dataset (see Supplementary Methods S7).

### Estimate of genetic parameters

For each of our fitness metrics, we report the additive genetic variance V_A_ and the narrow-sense heritability, h^2^ (Hendry *et al*. 2018; Bonnet *et al*. 2019a), on both the liability and the observed scales (de Villemereuil *et al*. 2016); see Supplementary Methods S4, “Glossary”. Narrow-sense heritability was calculated as V_A_/V_P_ (additive genetic variance divided by total phenotypic variance). On the observed scale, h^2^ was adjusted to account for the fixed effects using the QGglmm R package; accounting for fixed effects is not straightforward on the liability scale (de Villemereuil *et al*. 2018). Genetic correlations (r*_a_*) were calculated as: 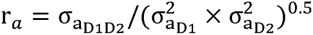. Phenotypic correlations (r*_p_*) were calculated as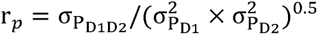, where 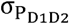 represents the total phenotypic covariance and 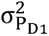 and 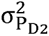 represent the total phenotypic variances of the two fitness metrics.

Finally, we obtained an estimate of V_A_(*w*) (additive genetic variance of relative fitness) by back-transforming the liability-scale estimates using the protocol detailed in Bonnet *et al*. (2022) (Supplementary Results S8-S9). For the heritability of relative individual fitness, we divided V_A_(*w*) by the total variance in relative fitness (Lynch & Walsh 1998).

## RESULTS

### Univariate analyses of fitness metrics

Among the time-limited fitness metrics, we found measurable, non-zero V_A_ for annual survival, annual fitness, and triannual breeding success, but not for annual breeding success (Figure 1; Table 2). Annual survival showed the highest heritability among time-limited fitness metrics (h^2^_observed_=0.2317, Table 3). Annual fitness and triannual breeding success were weakly heritable in our data set (respectively h^2^_liability_=0.0189 and h^2^_liability_=0.0807; Table 3).

**Figure 1.**
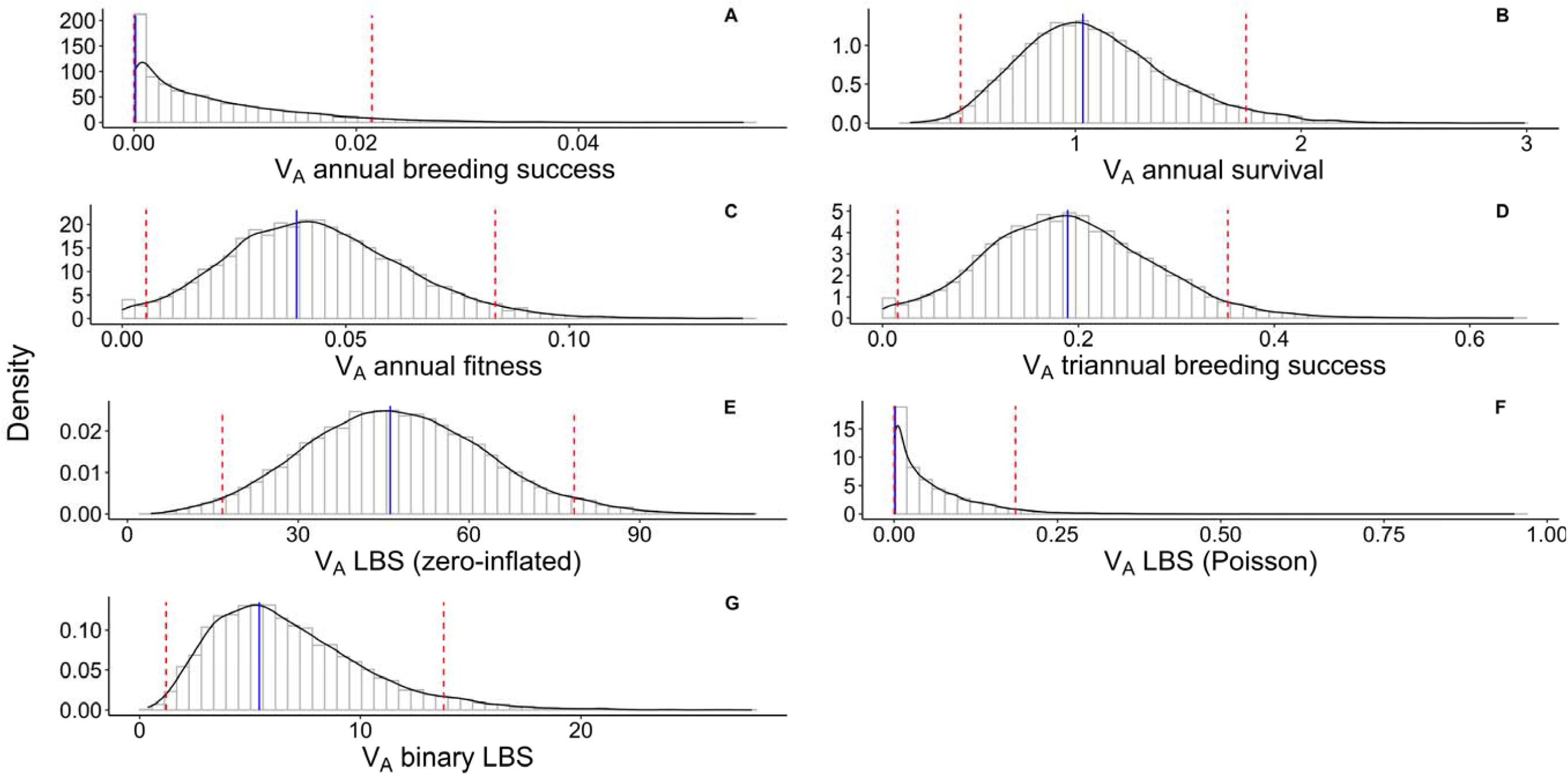
V_A_ on the liability scale of (A) *annual breeding success*, (B) *annual survival*, (C) *annual fitness*, (D) *triannual breeding success*, (E) *LBS* (zero-inflated component), (F) *LBS* (Poisson component), (G) *binary LBS*; estimated with univariate animal models and represented by their posterior MCMC distribution (bars), overlapped with their kernel density (solid black line). 95% credible intervals are indicated by red dashed lines; posterior mode is indicated by solid blue line.

**Table 2.** Univariate animal models: posterior modes (and 95% credible intervals, CI) of all variances associated to random effects for all different fitness metrics. Variances are on the latent scale and are marked in **bold** when the lower bound of the 95% credible interval is > 0.001^a^. Values below 10^-4^ are marked 0.

| **Trait** | **Variable** | **Posterior mode of the variance** | **95% CI** |
| --- | --- | --- | --- |
| Annual breeding success | Additive genetic effect ($V_A$) | 0.0001 | [0, 0.0214] |
|  | Permanent environment | 0 | [0, 0.0088] |
|  | Maternal effect | 0.0001 | [0, 0.0131] |
|  | **Hydroyear** | **0.0262** | **[0.0019, 0.0641]** |
|  | **Group-hydroyear** | **0.0323** | **[0.0051, 0.0766]** |
| Annual survival | **Additive genetic effect ($V_A$)** | **1.0336** | **[0.4913, 1.7559]** |
|  | Permanent environment | 0.0005 | [0, 0.1906] |
|  | Maternal effect | 0.0004 | [0, 0.1580] |
|  | Hydroyear | 0.0002 | [0, 0.0843] |
|  | **Group-hydroyear** | **0.1455** | **[0.0595, 0.2541]** |
| Annual fitness | **Additive genetic effect ($V_A$)** | **0.0390** | **[0.0053, 0.0834]** |
|  | Permanent environment | 0.0001 | [0, 0.0129] |
|  | Maternal effect | 0.0001 | [0, 0.0193] |
|  | **Hydroyear** | **0.0487** | **[0.0131, 0.1011]** |
|  | **Group-hydroyear** | **0.0744** | **[0.0322, 0.1243]** |
| Triannual breeding success | **Additive genetic effect ($V_A$)** | **0.0189** | **[0.0157, 0.3527]** |
|  | Permanent environment | 0.0006 | [0, 0.1063] |
|  | Maternal effect | 0.0009 | [0, 0.1533] |
| LBS, zero-inflated component | **Additive genetic effect ($V_A$)** | **46.1429** | **[16.6974, 78.4829]** |
|  | Maternal effect | 0.1286 | [0, 16.9133] |
|  | Hydroyear of birth | 0.0084 | [0, 5.9133] |
| LBS, Poisson component | Additive genetic effect | 0.0015 | [0, 0.1858] |
|  | Maternal effect | 0.0005 | [0, 0.1407] |
|  | Hydroyear of birth | 0.0009 | [0, 0.1227] |
| Binary LBS | **Additive genetic effect ($V_A$)** | **5.4168** | **[1.1961, 13.7798]** |
|  | Maternal effect | 0.0039 | [0, 2.1083] |
|  | Hydroyear of birth | 0.0069 | [0, 0.8008] |

<sup>a</sup> Note that since these are variances estimates, they cannot be lower than 0.

**Table 3.** Univariate animal models: posterior modes (and 95% credible intervals, CI) of heritability of different fitness metrics. Fitness metrics are marked in **bold** when the lower bound for the 95% credible interval for additive genetic variance (V_A_) > 0.001 (on the liability scale) (see Table 2). Values below 10^-4^ are marked 0.

| Fitness metric | $h^2$ on the liability scale<sup>a</sup> | | $h^2$ on the observed scale | |
| --- | --- | --- | --- | --- |
|  | Posterior mode | 95% CI | Posterior mode | 95% CI |
| Annual breeding success | 0.0001 | [0, 0.0196] | 0 | [0, 0.0048] |
| **Annual survival** | **0.4485** | **[0.3908, 0.5837]** | **0.2317** | **[0.1498, 0.3309]** |
| **Annual fitness<sup>b</sup>** | **0.0189** | **[0.0024, 0.0376]** | «0» 0.0051 | [0, 0.0094] |
|  |  |  | «1» 0.0003 | [0, 0.0008] |
|  |  |  | «2» **0.0064** | **[0.0010, 0.0131]** |
| **Triannual breeding success<sup>b</sup>** | **0.0807** | **[0.0156, 0.1543]** | «0» **0.0134** | **[0.0014, 0.0247]** |
|  |  |  | «1» 0.0011 | [0, 0.0031] |
|  |  |  | «2» **0.0134** | **[0.0025, 0.0254]** |
|  |  |  | «3» 0.0012 | [0, 0.0029] |
| LBS |  |  |  |  |
| • Zero-inflated component | **0.8928** | **[0.6062, 0.9460]** | **0.5097** | **[0.3605, 0.5674]** |
| • Poisson component | 0.0019 | [0, 0.4466] | 0.0007 | [0, 0.1311] |
| **Total** | NA<sup>c</sup> | NA | **0.2511** | **[0.1768, 0.3353]** |
| **Binary LBS** | **0.8385** | **[0.5593, 0.9289]** | **0.4480** | **[0.3114, 0.5167]** |

<sup>a</sup> To estimate heritability on the liability scale we added the link variance to the total phenotypic variance ( $V_P$ ).
<sup>b</sup> For annual fitness and triannual breeding success, a separate value of $h^2$ on the observed scale was produced for each ordinal value of the trait («ordinal value» in the table); this occurs when parameters of ordinal (multiple threshold) models are back-transformed to the observed scale.
<sup>c</sup> Total $h^2$ on the liability scale is not provided because it is not easily interpretable for zero-inflated Poisson models.

We also found high additive genetic variance for lifetime fitness in female Amboseli baboons (Figure 1, Table 2). In particular, heritabilities for binary LBS and the zero-inflated component of LBS were high and quite similar (as expected, given that they capture a similar biological phenomenon–survival to reproductive age): h^2^_observed_=0.4480 for binary LBS and h^2^_observed_=0.5097 for the zero-inflated component of LBS (Table 3). In contrast, the CI for estimates of V_A_ for the Poisson component of *LBS*, which reflects the count of surviving offspring, overlapped 0 (Table 2).

Consequently, the heritability for LBS overall was intermediate between the heritabilities of its two components (h^2^_observed_=0.2511, Table 3). Following Bonnet *et al*. (2022) we also obtained an estimate of V_A_(*w*) (additive genetic variance for relative LBS on the observed scale) that was also very high (V_A_(*w*)=0.9023). Its heritability (h^2^_observed_=0.4701) was comparable to the heritability of the zero-inflated component of LBS. For a discussion of this value in comparison with that obtained by Bonnet et al., 2022, V_A_(*w*)=0.231, see Supplementary Results S8.

Considering other random effects, we found hydroyear effects for annual breeding success and annual fitness, indicating non-genetic variance associated with random year-to-year variation in the environment (Figure 2; Table 2). We also found effects of group-hydroyear for all time-limited fitness metrics, indicating non-genetic variance associated with differences among social groups specific to different hydroyears (Figure 2; Table 2). In contrast, 95% CIs overlapped zero for permanent environment and maternal effects in the annual and triannual models (Table 2). For the influence of fixed effects and non-genetic sources of variance on fitness metrics, see Supplementary Results S10, Table S3, and Supplementary Figures S1-S2.

**Figure 2.**
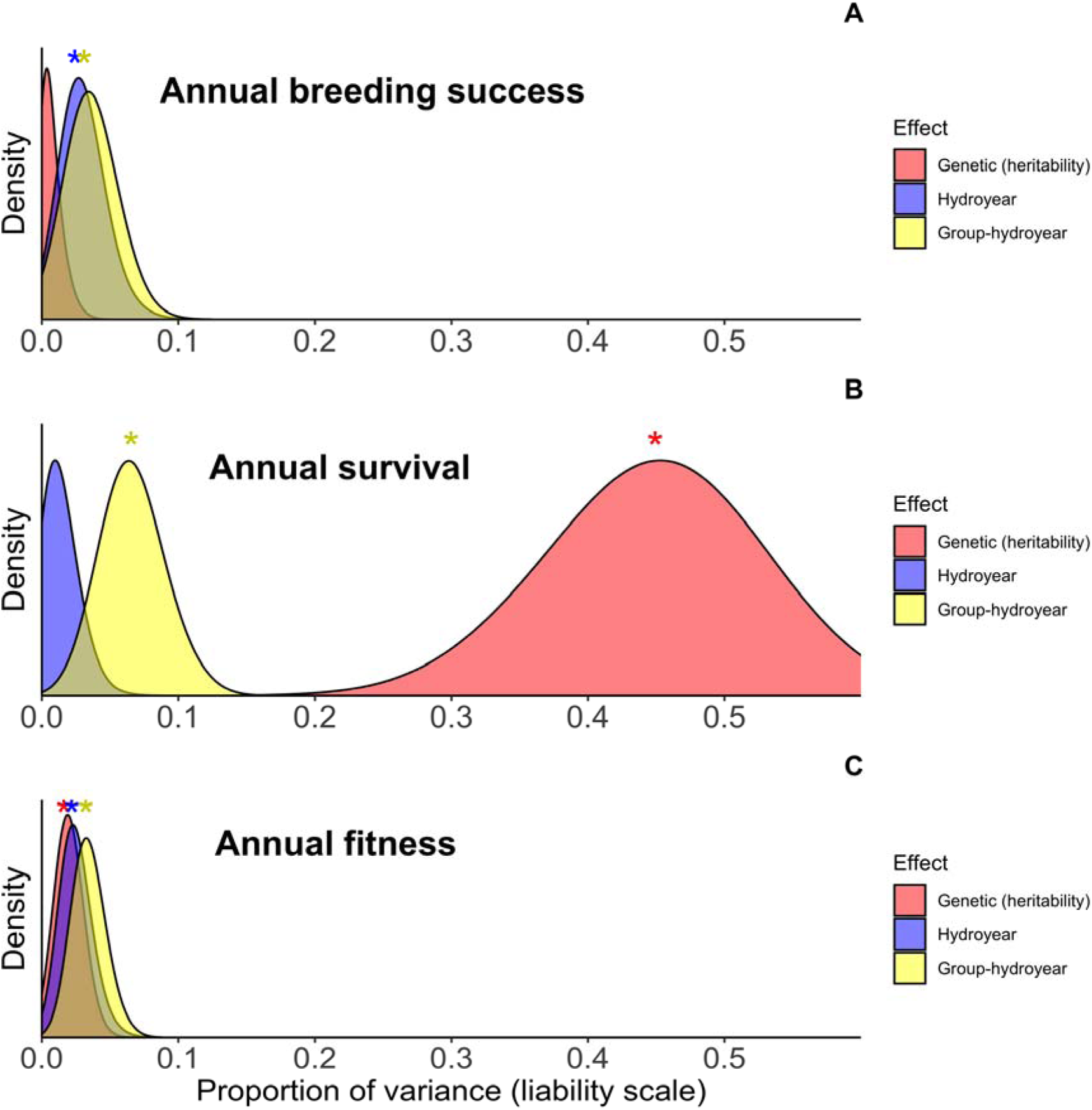
Variance components (proportion of phenotypic variance) for (A) *annual breeding success*, (B) *annual survival*, (C) *annual fitness*, estimated with univariate animal models and represented by the density of their posterior MCMC distribution on the liability scale. Heritability (h^2^) is shown in red, hydroyear effects in blue, and group-hydroyear effects in yellow: asterisk indicates that the lower bound of the 95% credible interval is greater than 0.001. Permanent environment and maternal effects are not presented as the posterior density was heavily weighted towards zero. Distributions are scaled to aid comparisons.

### Bivariate analysis of fitness metrics

We found very high positive genetic correlations (r*_a_*) between all fitness metrics (Table 4). Genetic correlations between the three annual metrics, and between the annual metrics and triannual breeding success, were particularly high (r*_a_*>0.98; Table 4). Genetic correlations were also high between the time-limited fitness metrics and binary LBS (range r*_a_*=0.89–0.99). The correlation between annual breeding success and binary LBS was the lowest of this set, albeit still substantial (r*_a_*=0.89), and all correlations with annual breeding success produced relatively wide credible intervals (Table 4). Notably, the credible intervals showed some overlap for all the genetic correlations between the time-limited fitness metrics and binary LBS, suggesting that they capture similar information. Genetic correlations between time-limited fitness metrics and LBS were weaker in comparison (despite being still relatively high; Table 4). They ranged from r*_a_*=0.8837 between LBS and annual fitness to r_a_=0.7299 between LBS and annual breeding success (Table 4). Correlations on the latent scale were very similar to those obtained on the observed scale. Models run with permuted datasets showed much lower, non-significant genetic correlations, indicating that these results were not driven by the model structure and implementation (see Supplementary Methods S7).

**Table 4.** Bivariate analyses of fitness metrics. Posterior modes and 95% credible intervals of correlations between fitness metrics on the observed scales. Additive genetic correlations are placed above the diagonal (boxes are shaded light gray); phenotypic correlations are placed below the diagonal (boxes are shaded dark gray). Correlations marked in **bold** are different than 0 (credible intervals do not contain zero).

|  | Annual<br>breeding success | Annual survival | Annual fitness | Triannual<br>breeding success | LBS | Binary LBS |
| --- | --- | --- | --- | --- | --- | --- |
| Annual breeding<br>success |  | **0.9866**<br>[0.8228, 0.9974] | **0.9854**<br>[0.9201, 0.9996] | **0.9854**<br>[0.8426, 0.9992] | **0.7299**<br>[0.6134, 0.8829] | **0.8908**<br>[0.2090, 0.9988] |
| Annual survival | **0.0450**<br>[0.0187, 0.0599] |  | **0.9971**<br>[0.9936, 0.9999] | **0.9977**<br>[0.9030, 0.9997] | **0.8529**<br>[0.7527, 0.9166] | **0.9955**<br>[0.9757, 0.9997] |
| Annual fitness | **0.1292**<br>[0.1195, 0.1361] | **0.2508**<br>[0.2212, 0.2827] |  | **0.9938**<br>[0.9622, 0.9997] | **0.8837**<br>[0.8302, 0.9435] | **0.9791**<br>[0.9087, 0.9970] |
| Triannual<br>breeding success | 0.0011<br>(-0.0043, 0.0095] | **0.0768**<br>[0.0489, 0.1028] | **0.0456**<br>[0.0349, 0.0550] |  | **0.7884**<br>[0.6795, 0.8641] | **0.9626**<br>[0.6236; 0.9971] |
| LBS | **0.0320**<br>[0.0166, 0.1095] | **0.1455**<br>[0.0914, 0.1822] | **0.1321**<br>[0.0934, 0.1538] | **0.1754**<br>[0.1150, 0.2419] |  | **0.9822**<br>[0.9547; 0.9969] |
| Binary LBS | **0.0201**<br>[0.0031, 0.0448] | **0.3366**<br>[0.2934, 0.3803] | **0.0767**<br>[0.0567, 0.1011] | **0.0680**<br>[0.0301, 0.0971] | **0.4984**<br>[0.4351, 0.5477] |  |

Phenotypic correlations (r*_p_*) were all positive but lower and more variable. The highest phenotypic correlations were between LBS and binary LBS (r*_p_*=0.4984), annual survival and binary LBS (r*_p_*=0.3366), annual fitness and annual survival (r*_p_*=0.2508), and triannual breeding success and LBS (r*_p_*=0.1754). All other phenotypic correlations were <0.15, although only the correlation between annual and triannual breeding success overlapped zero (Table 4). For the bivariate model estimates of heritability and non-genetic variances see Supplementary Results S11, and Supplementary Table S4.

## DISCUSSION

Our results show that genotype plays a substantial role in explaining variance in lifetime fitness and annual survival in female Amboseli baboons. We also show that the bulk of the genetic contribution to survival occurs during the infant and juvenile stages, prior to first reproduction, reflecting variation in the ability to survive through the infant and juvenile period. Thus, as baboons mature, a shift occurs in the relative contribution of genes versus the environment to fitness. However, all our fitness metrics exhibited high positive genetic correlations, indicating that, whether fitness is measured over the full life course or on shorter time scales, it involves a consistent underlying genetic architecture. Below, we discuss both the evolutionary and methodological implications of these findings for understanding evolution in natural populations, including the surprisingly high levels of additive genetic variance for several of the fitness measures we considered—a result that contradicts expectations for populations at equilibrium.

### The genetic components of fitness metrics in female Amboseli baboons

Our results identify survival to the first production of a surviving offspring as the life history trait for which genetic differences among females are most influential. In contrast, the total number of offspring produced beyond the first is much less affected by genotype and is better explained by environmental and stochastic effects on female fertility in her reproductive years. These conclusions stem from the observation that the additive genetic variance (V_A_) of lifetime breeding success (LBS) was entirely associated with the zero-inflated component of the LBS distribution, which essentially corresponds to survival to the first successful reproduction (Moiron *et al*. 2022). In contrast, the number of infants produced after first reproduction (Poisson component) did not appear to possess measurable V_A_ (Tables 2, 3). Accordingly, while the genetic correlation between the LBS and binary LBS – i.e. survival to first reproduction – indicates an almost complete genetic correspondence (r_a_=0.9822), the phenotypic correlation is much lower (r*_p_*=0.4984), indicating that binary LBS captures all the genetic variance of LBS, but it does not capture all the variance in LBS that is not genetic. In other words, the additive genetic effects on binary LBS and LBS are nearly identical, but the environmental effects on binary LBS and LBS differ in kind, magnitude, or both. Accordingly, the heritability of binary LBS (h^2^_observed_=0.4480) is higher than that of LBS, because while the two metrics share the same genetic variance, LBS exhibits greater phenotypic variance than binary LBS, as it includes the non-genetic variances of both zero-inflated and Poisson components.

Correspondingly, among the time-limited fitness metrics, annual survival is relatively highly heritable (h^2^_observed_=0.2317; Table 3). However, annual breeding success does not appear to harbor appreciable genetic variance, a result in line with some other studies of time-limited fitness metrics (McFarlane *et al*. 2014; Wolak *et al*. 2018; Moiron *et al*. 2022). That is, whether a female baboon can produce a live offspring consistently on an annual basis does not appear to be measurably influenced by an individual’s genotype, but only by environmental and social factors (partially captured by hydroyear and group-hydroyear: see Supplementary Results S10). The heritability of triannual breeding success, i.e. breeding success measured over three-year intervals, was low but bounded away from zero (Table 3), reflecting the fact that for long-lived, non-seasonal reproducers, like baboons, differences in the rate of offspring production might be better captured by longer time intervals, which allow variation in breeding success to emerge over fewer total intervals. Overall, the low heritability of time-limited metrics of breeding success aligns with the lack of V_A_ in the Poisson component of LBS, as both indicate that the realized rate of infant production is less influenced by genotype than survival (especially survival to first reproduction).

### Implications of the high V_A_ and h^2^ of fitness in our population

The heritability of LBS in female baboons in Amboseli (h^2^_observed_=0.2511) is high for a lifetime fitness metric (the estimated mean for wild and semi-wild populations of birds and non-human mammals < 0.1; Hendry *et al*. (2018); Bonnet *et al*. (2022)). Fitness measured in other long-term studies of animal populations sometimes reveals measurable additive genetic variance, but heritability is usually much smaller (Hendry *et al*. 2018). The additive genetic variance in fitness corresponds to the upper limit of phenotypic change that can happen in a population: females in this population thus show considerable genetic evolutionary potential. However, high V_A_ does not necessarily imply the same potential for phenotypic change, even when a trait is under selection (Gauzere *et al*. 2022; Pemberton *et al*. 2022) or if the trait is fitness itself (Bonnet *et al*. 2022). Directional selection in wild populations frequently does not lead to a corresponding phenotypic change (“paradox of stasis”, Pujol *et al*. (2018). The high heritability of fitness implies, for example, that mean absolute fitness (i.e., LBS) in this population has the genetic capacity to increase over time. If the V_A_ and heritability we report for LBS were extrapolated to a predicted phenotypic response to selection, population sizes should have grown exponentially: based on our estimates, mean LBS is expected to have quintupled in two generations, something that is clearly not occurring in the Amboseli baboons, where the population has grown modestly since the 1970’s (Supplementary Results S9; Bonnet *et al*. (2022). This contradiction raises the important question of why female baboons in this population have not evolved to maximize their fitness, in spite of the genetic potential to do so.

One explanation for this apparent stasis is environmental change. If the environment deteriorates, organisms may experience phenotypic plasticity (non-genetic) that translates to lowered LBS, despite selection for higher LBS. Under this scenario, environmentally-induced changes in phenotype can overshadow genetic evolution, so that genetic change is counterbalanced by environmental shifts (Bonnet *et al*. 2017; Hunter *et al*. 2022). This kind of “cryptic evolution” due to environmental deterioration has been found to mask evolutionary change in other long-term studies, (Merilä *et al*. 2001; Wilson *et al*. 2007). In Amboseli, alleles that predispose to higher LBS might be selected, even while changes in the environment impeded an observed increase in the LBS phenotype itself. The Amboseli ecosystem is a highly dynamic environment with considerable rainfall variability and fluctuating populations of plants, herbivores, and predator communities, including dramatic declines over time in key plant species for baboons, such as *Vachellia xanthophloea* (Western & Van Praet 1973; Western & Behrensmeyer 2009; Alberts & Altmann 2012; Okello *et al*. 2016). Since three quarters of the variance in LBS is non-genetic (Table 2), even relatively small detrimental changes in the environment could mask phenotypic evolution.

As other fitness metrics are highly genetically correlated with LBS, we examined the possibility that annual survival (the short-term metric with the highest heritability) could indeed be evolving (sensu Bonnet *et al*. (2017); (Bonnet *et al*. 2019b; Gauzere *et al*. 2022). In support of this idea, the estimated breeding values (EBVs) for annual survival increased at a rate of 0.013 [0.006, 0.022] per hydroyear over the course of the study (see Supplementary Results S12, Supplementary Figures S3-S4; Hadfield *et al*. (2010); Bonnet *et al*. (2019b); Tuliozi *et al*. (2023). At face value, this change is predicted to translate into a 14.8% higher probability of annual survival for an individual born in 2022 compared to one born in 1971 (Supplementary Figure S4), but no such change is readily apparent at the phenotypic level. Nevertheless, the increase in EBVs that we observed for annual survival is consistent with a successful evolutionary response to directional selection, which is phenotypically masked by counteracting changes in the environment.

Together, our findings raise the question of how substantial V_A_ for fitness is maintained, since populations at equilibrium are predicted to have low V_A_ for fitness. Fluctuating selection in dynamic ecosystems can preserve overall genetic variation (Bergland *et al*. 2014; Bitter *et al*. 2024), but the high V_A_ for fitness and increasing EBVs for annual survival in the study population suggest that the relationship between genotype and fitness remained relatively consistent during our study period. Fluctuating selection could still be acting on longer timescales in Amboseli, but is unlikely to explain the results we present here (see Supplementary Results S12).

An additional explanation for the large V_A_ for fitness is that indirect effects linked to the social environment may have inflated our V_A_ and heritability estimate for (binary) LBS. Indirect effects refer to the influence that individuals have on each other’s phenotypes, and are frequently cited as an important, yet underexplored, explanation for apparent evolutionary stasis (Pujol *et al*. 2018; Fisher & McAdam 2019; McGlothlin & Fisher 2021). In the Amboseli baboon population, the death of the mother during the first year of an infant’s life is usually fatal to the infant, and the death of the mother during the first four years of a female’s life can reduce her lifespan even if she survives to adulthood (Tung *et al*. 2023). Because maternal survival is correlated with offspring survival and mothers share genes with their offspring, V_A_ estimates will be inflated if the correlation stems from non-genetic covariance but is attributed to genotype. Moreover, high maternal social rank is associated with increased infant survival in the study population (Silk *et al*. 2003; Creighton *et al*. 2025), and female rank in cercopithecine primates is socially inherited along matrilines. As a result of this non-genetic “inheritance” of a trait that influences infant survival, the fitness consequences of variance in social status might be misattributed to genetic variation. While modeling indirect effects (both genetic and non-genetic) falls outside of the scope of this study, quantifying indirect genetic effects on fitness and disentangling social and genetic effects remains an important frontier for studies of contemporary evolution in natural populations (Young *et al*. 2019).

Finally, our estimate of the evolutionary potential of the population could change if we included males in our analysis. The challenges that females and males experience in this population are very different: philopatric females remain with their female relatives throughout life while males disperse to find different groups, where they fight to establish rank and gain access to females. Given the divergence in male and female life history, genotypes that are beneficial for females might not be beneficial for males, creating negative genetic covariance between the genetic components of female and male LBS (i.e. sexual antagonistic pleiotropy, Brommer *et al*. (2007b). Alternatively, even in the absence of sexually antagonistic pleiotropy, if V_A_ for fitness was lower in males than in females, the genetic variance for fitness in both sexes would be reduced, a prediction consistent with previous, lower estimates of V_A_(*w*) when males and females were considered together (Bonnet *et al*. (2022): Supplementary Results S8). Understanding differences between the sexes in the genetic component of fitness—whether due to changes in sign or magnitude—is another important goal for assessing the potential for responses to natural selection in the wild (Wolak *et al*. 2018).

### Genetic correlations between fitness metrics

We identified universally high genetic correlations between all the metrics we studied. These include measures that focus on survival versus those that focus on fertility, as well as those that consider time-limited versus lifetime fitness metrics. Our findings reinforce the growing consensus that survival and reproduction often do not exhibit apparent genetic tradeoffs (Chang *et al*. 2024). We caution, however, that because survival has a causal effect on reproductive success (i.e., reproduction ceases with death, Tuliozi *et al*. (2022), a subtle trade-off could still be present but masked.

Our finding that time-limited and lifetime fitness metrics are strongly genetically correlated in female baboons (Table 4), despite having very different heritability values, has also potential practical implications. Our results suggest that, for quantitative genetics purposes, time-limited metrics are an accurate proxy of lifetime breeding success and may still provide information on the genetic basis of individual fitness when LBS is not available. In the Amboseli baboons, a metric based on the first successful reproduction (binary LBS) would serve as a similarly reliable proxy that is measurable on a much shorter timescale. The degree to which this result generalizes to other systems remains an important question, but we speculate that it is most likely to hold in long-lived species with slow development.

Notably, longevity is the main contributor to variation in phenotypic fitness in long-lived female mammals: variation in how long females live is much more important for female lifetime reproductive success than variation in the relative frequency of offspring production. Here, we show that both annual and triannual breeding success are nevertheless genetically correlated with LBS and annual survival, likely because living longer results in more opportunities to reproduce, and thus more offspring. By itself, however, consistency in the annual production of offspring is not measurably heritable. This incongruency may explain why genetic correlations that involve annual breeding success tend to show wider credible intervals than genetic correlations among the other fitness metrics (Table 4).

Finally, the positive genetic correlations between reproduction and survival in our study population indicate that female baboons do not experience a genetic tradeoff between the two fitness components. Interestingly, a phenotypic trade-off between adult female reproduction and survival was identified in this population after adjusting for female phenotypic quality (McLean *et al*. 2019). In light of the genetic correlations we report here, this phenotypic tradeoff is likely to be unlinked to genotype, and possibly accounted for by the finite resources available to adult females (Van Noordwijk & De Jong 1986). Together, these observations underscore the importance of investigating sources of variance in fitness at both the genetic and the phenotypic levels, as they provide distinct but complementary information on how natural populations evolve.

## Author contributions

B.T.: conceptualization, methodology, formal analysis, writing (original draft and review/editing). E.A.A.: resources, data curation, writing (review/editing) and funding acquisition. J.T.: resources, data curation, writing (review/editing), and funding acquisition. S.C.A.: conceptualization, resources, data curation, writing (review/editing) and funding acquisition.

## Acknowledgements

We thank Duke University, Princeton University, the University of Notre Dame, the Max Planck Institute for Evolutionary Anthropology for financial and logistical support at various times over the years. For assistance and cooperation in Kenya, we are grateful to the Kenya Wildlife Service (KWS), the Wildlife Research Training Institute (WRTI), University of Nairobi, the Kenya Institute of Primate Research (IPR), National Museums of Kenya, National Environment Management Authority, and National Commission for Science, Technology, and Innovation (NACOSTI). We also thank the members of the Amboseli-Longido pastoralist communities, the Enduimet Wildlife Management Area, Tortilis Camp, Ker & Downey Safaris, Serena Lodge, Air Kenya, and Safarilink. Particular thanks go to the current Amboseli Baboon Project long-term data collection team (J.K. Warutere, L. Musembei, D.K. Tajeu), and to T. Wango, and V. Oudu for their assistance in Nairobi. The baboon project database, Babase, is managed by J. Gordon and C. Broderick. For a complete set of acknowledgments please visit http://amboselibaboons.nd.edu/acknowledgements/.

