## Supplemental Information for "Lifetime fitness and annual survival are heritable and highly genetically correlated in a wild primate population"

SUPPLEMENTARY INFORMATION

**Summary**

Supplementary Methods

**S1**. Fixed effects……..……………………………………………………………..…………………2

**S2**. Genetic ancestry…………………………………………………………………………………..5

**S3**. Background relatedness and missing pedigree links………………………….…………..………6

**S4**. Glossary……………………………………………………..…….………………………………7

**S5**. Variance-covariance random structure of bivariate models……………………………………….9

**S6**. Full list of random effects structures of bivariate models………………………………………..10

**S7**. Permutation of bivariate analysis…………………………………………………………..…….12

Supplementary Results

**S8**. V_A_(*w*) estimates obtained in Amboseli baboons……………………………………………..…..13

**S9**. Expected increase in relative fitness in Amboseli baboons………………………………….…..15

**S10**. Non-genetic sources of variance and fixed effects……………………………………………..15

**S11**. Heritability of fitness metrics estimated from bivariate models………………………………..16

**S12**. Evolutionary change in annual survival………………………………………………………...17

Supplementary Tables

**Table** **S1**. Univariate animal models including genetic ancestry scores: posterior modes of all fixed effects. ……………………………………………………..………………………………….……..19

**Table** **S2**. Model parameters for both univariate and bivariate animal models. ……….….…….…..21

**Table** **S3**. Univariate animal models: posterior modes of all fixed effects………………………….22

**Table** **S4**. Bivariate animal models: posterior modes for all observed heritabilities obtained from bivariate models of fitness metrics…………………………………………………………………..24

Supplementary figures

**Figure S1**. Average annual survival as a function of age class in female baboons…………….…….26

**Figure S2**. Average annual breeding success as a function of age class in female baboons…………27

**Figure S3**. Distribution of regression coefficients (slopes) for change in estimated breeding values of annual survival………………………………………………………………………………………..28

**Figure S4**. Genetic trend for annual survival…………………………………………………………29

Supplementary Methods

### **S1. Fixed effects**

Models for annual breeding success, annual survival, and annual fitness were fitted with the same fixed effects. All fixed effects are included in both univariate and bivariate models as presented.

i) **Age**: continuous covariate.

Definition: For each female-hydroyear, ‘age’ was defined as the age of the female on the birthday that fell within that hydroyear, plus one (i.e., the period between 0 and 1 is considered the first year of life). For example, the female AKP, born on the 1^st^ of August 2015, was assigned an age of 3 during the female-hydroyear AKP-2017, which extended from 1^st^ of November 2016 to the 31^st^ of October 2017.

Rationale: An individual’s age is known to strongly affect time-limited metrics of fitness in most organisms, and in the Amboseli baboons in particular, both because juveniles do not reproduce and because reproductive success and survival strongly follow age-linked patterns (Jones *et al.* 2014; Campos *et al.* 2020).

ii) **Age squared**: continuous covariate.

Definition: Age raised to the power of two.

Rationale: This effect captures non-linear effects of age on reproductive success and survival. In our study population the average reproductive success is very low for the first four to six years of females’ lives and peaks for females between eight and eleven years old. It then begins decreasing around the age of sixteen (Alberts & Altmann 2003; Altmann & Alberts 2003a, b; Charpentier *et al.* 2008; Colchero *et al.* 2021).

iii) **Group size**: continuous covariate.

Definition: The number of sexually mature females in the individual’s group, averaged across the hydroyear. In hydroyears with permanent group fissions or fusions, we used the average group size of the group in which the female spent the majority of her time. Mean group size of study groups between 1971 and 2022 was 15.64 ± 5.86 s.d. adult females.

Rationale: Physiology, behavior, fertility and survival all vary with group size in the Amboseli baboons (Altman & Alberts 2003; Charpentier *et al.* 2008; Lea *et al.* 2015; Markham *et al.* 2015).

iv) **Group size squared**: continuous covariate.

Definition: Group size raised to the power of two.

Rationale: This effect captures possible non-linear influences of group size on annual fitness metrics. For example, previous analyses have suggested advantages associated with intermediate group size (Markham *et al.* 2015).

v) **Habitat quality**: binary effect.

Definition: We fitted habitat quality as a binary fixed effect, assigning each female-hydroyear to either a low- or high-quality habitat depending on where the individual had spent the majority of a hydroyear.

Rationale: The two original study groups lived in a low-quality habitat until the late 1980s, where acacia trees were scarce. One study group shifted its home range in 1987 approximately 5-6 km southwest of the original home range to an area with abundant trees; the second shifted its home range to the same area in 1991 (Gesquiere *et al.* 2018). All study groups have since resided in the higher quality habitat, and females of the population have since experienced shorter interbirth intervals and increased offspring survival (Altman & Alberts 2003; Alberts *et al.* 2005; Gesquiere *et al.* 2018; Anderson *et al.* 2024).

vi) **Rainfall anomaly of the previous hydroyear**: continuous covariate.

Definition: Difference between total rainfall in a given hydroyear and the average total rainfall calculated over all the hydroyears during the entire study period. Rainfall data have been collected by the research project since 1976. For rainfall information before that date, we employed gridded datasets provided by the Global Precipitation Climatology Centre (Schneider *et al.* 2008).

Rationale: Rainfall is linked to differences in survival (Campos *et al.* 2017; Campos *et al.* 2021) and fertility (Lea *et al.* 2015). We used rainfall anomaly of the previous hydroyear because we hypothesized that this value could impact resource availability – and thus female condition – during key times such as conception and pregnancy.

While annual metrics refer to hydroyears and thus have the same timeframe for all individuals, triannual breeding success was individually calculated, starting on the day of each female’s birth. Therefore, while annual metrics were based on demographic events that occurred within each hydroyear, triannual breeding success was based on demographic events that occurred within three-year intervals (triennia) where dates were specific to each female. One consequence of this individual-specific timeframe was that we could not include hydroyear-linked non-genetic sources of variance, such as hydroyear and group-hydroyear, because each three-year interval started and ended in different dates. However, we believe that this decision is unlikely to have substantially influenced the estimates of the genetic variance of triannual breeding success. When using unaligned timeframes as records, yearly differences in the physical and social environment should not show bias across the dataset, or covary with the genetic component. In other words, unaligned timeframes might increase residual variance, but without systematically increasing additive genetic variance. Analyses modeling triannual breeding success included the following fixed effects both in univariate and bivariate models (the biological rationales are the same as for the corresponding fixed effects in the annual models):

i) **Triennium**: continuous covariate.

Definition: Starting from birth, an individual lifetime was divided in three-year long intervals, which were numbered starting from the earliest (1^st^) triennium, from birth to 3^rd^ birthday. Analogous to age in the annual models.

ii) **Triennium squared**: continuous covariate.

Definition: Triennium raised to the power of two.

iii) **Triennium group size**: continuous covariate.

Definition: The number of sexually mature females in the individual’s group, averaged for each female across each triennium of her life.

iv) **Triennium group size squared**: continuous covariate.

Definition: Triennium group size raised to the power of two.

v) **Habitat quality**: binary effect.

Definition: We assigned each female-triennium to high or low habitat quality depending on where the individual had spent the majority of the triennium.

vi) **Triennium rainfall anomaly**: continuous covariate.

Definition: The difference between the total cumulative rainfall in a three-year interval starting one year (365 days) before the beginning of a female-triennium and ending two years into the female-triennium, with the mean rainfall calculated over all possible three-year intervals. The shift of this window to one year before the start of the female-triennium parallels the window shift for hydroyear-based analyses. Here, it also reflects the delayed effects of low (or high) rainfall on ecosystem productivity, so a presumably lagged effect on female survival and/or reproduction.

Analyses modeling *LBS* and *Binary LBS* included the following fixed effects both in univariate and bivariate models (the biological rationales are the same for the corresponding fixed effects in the annual models):

i) **Rainfall anomaly a year before birth**: continuous covariate

Definition: We calculated rainfall anomaly as the difference between the cumulative rainfall in the one-year period preceding the individual’s birth, and the average rainfall of all one-year intervals.

ii) **Habitat quality at birth**: binary effect.

Definition: Individuals were either born in the low- or high-quality habitat.

iii) **Hydroyear of birth**: linear covariate.

Definition: Using the first hydroyear (cohort) in the dataset as a starting point (cohort 1), we assigned an increasing value to each hydroyear of birth.

Rationale: This effect can account for chronological linear changes in the environment and population.

### **S2. Genetic ancestry**

The population of baboons living in the Amboseli basin are an admixed population with majority yellow baboon (*Papio cynocephalus*) ancestry and minority anubis baboon ancestry (*P. anubis*). These two species are not strongly ecologically differentiated (Winder 2015; Wango *et al.* 2019) and produce viable and fertile offspring, although signatures of selection against admixture are detectable at the genomic level (Ackermann *et al.* 2006; Tung *et al.* 2008; Vilgalys *et al.* 2022). Anubis-yellow admixture has been linked to several traits with the potential to influence fitness in the Amboseli population, including maturation rates, mating success, and male-female affiliation (Charpentier *et al.* 2008; Tung *et al.* 2012; Fogel *et al.* 2021). We therefore explored whether genetic ancestry influences the fitness metrics we considered here by modeling genetic ancestry score – represented as the estimated proportion of introgressed anubis ancestry in each individual’s genome (Vilgalys *et al.* 2022) – as a fixed effect in our univariate models. Given that the subset of individuals for whom ancestry scores are available is currently considerably smaller than the full dataset (2644 female-hydroyears for 182 females, 79 females with full life history), we ran our models on the subset of the population for whom we had ancestry scores, both with and without ancestry score as a fixed effect, to check whether the genetic parameter estimates are affected by ancestry. Ancestry scores were z-transformed to facilitate comparison with other fixed effects.

We did not find an effect of genetic ancestry scores on any metric of fitness (Table S1). Because of the greatly reduced sample size in this analysis, the estimates for the variance parameters were mostly not recognizably different from zero and are not reported here. We can exclude that this outcome was caused by the inclusion of ancestry scores because when we ran the models with the same reduced sample size, but no ancestry score fitted as a fixed effect, we obtained variance estimates almost identical to models run with ancestry scores fitted as a fixed effect.

### **S3. Background relatedness and missing pedigree links.**

To estimate additive genetic variance, the animal model uses a matrix of genetic relationships (matrix **A**) derived from the pedigree. Thus, the accuracy of the pedigree (in terms of true levels of DNA shared identical-by-descent) will influence the accuracy of genetic estimates. The degree to which pedigrees are incomplete (e.g., due to missing paternities or maternities) or contain error (e.g., due to misassigned paternities) varies by population, but in general, is more likely to underestimate heritability than overestimate it because it underestimates resemblance between relatives (Cantet *et al.* (2000); Charmantier and Réale (2005); Morrissey *et al.* (2007); Mawass and Milot (2022); but see Wolak and Reid (2017) for an example of an overestimation of heritability). Furthermore, recent genomic work demonstrates that members of natural populations may exhibit appreciable “background relatedness”, i.e. low but widespread allele sharing between members that is not accounted for by recent pedigree relatedness (Freudiger *et al.* 2025). This discovery was made possible by measuring stretches of the genome inherited identical-by-descent between individuals, which reflects both recent and historical shared ancestry. To our knowledge, the potential effects of “background relatedness” on genetic variance estimates in wild populations are unknown.

A full comparison of heritability estimates obtained when using IBD relatedness (rIBD) versus relatedness obtained from the pedigree (rPed) to build the **A** matrix falls outside the scope of this study. However, to test if missing pedigree links and overall realized IBD relatedness was likely to decrease our heritability estimates, we re-ran our univariate models with a simulated additive genetic matrix that resembled an IBD relatedness matrix recently generated for a subset of baboons in the Amboseli (Galán Plana et al., *personal communication*). Because IBD estimates require genome-scale genotype data, only 9.2 – 9.3% of dyads in our main models also have corresponding estimates of IBD relatedness, precluding us from using the genomic data to independently calculate the heritability of fitness metrics (Bérénos *et al.* 2014).

Nonetheless, a comparison between the relatedness values obtained from the pedigree (rPed) and the rIBD values in the subset of the population for which both were available allowed us to simulate rIBD values for the additive genetic matrices obtained from our trimmed pedigrees. In other words, we simulated values of IBD relatedness consistent with those found in our population, and we put them in place of the current rPed values in our **A** matrices. We therefore adjusted the rPed values in our **A** matrices to make them consistent with rIBD values, as follows.

First, to simulate background relatedness and the potential for missing links, we added ≈ 0.1 (mean addition = 0.1058, range 0.100 – 0.705) to all relationships in the **A** matrix that had a rPed < 0.25; and we added ≈ 0.03 (mean addition = 0.0326, range 0.030 – 0.500) to all relationships in the **A** matrix that had a rPed >= 0.25. Since absolute rIBD values in the population have yet to be fully validated, we chose to simulate rIBD values in the top range of potential rIBD values – i.e., we preferred to simulate overestimated rIBD values to perform a conservative test of whether the true IBD relatedness would not impact our estimates. Due to issues with the matrix decomposition algorithm of MCMCglmm, we could substitute the simulated rIBD values only in the **A** matrices of univariate models (as these **A** matrixes were smaller and less complex). We therefore focused our test on the **A** matrix of the univariate binary LBS model. Finally, to make the **A** matrix with simulated rIBD positive definite, we added 0.5 to all diagonal values of the **A** matrix. This step precluded comparison with our original results (increasing diagonal values in an **A** matrix can decrease additive genetic variance), so we also re-ran our binary LBS model with a version of the original **A** matrix, unmodified except for the same addition of 0.5 to the diagonal values.

The binary LBS models run with simulated rIBD returned genetic variance and heritability estimates very similar to the values obtained by the corresponding models run with rPed (respectively, h^2^ = 0.30 and h^2^ = 0.33; credible intervals mostly overlap). Note that both these models had increased diagonal values, and yet they were fairly similar to the h^2^ obtained in the original binary LBS model (h^2^ = 0.43). Given that the simulated rIBD values are likely to be either equal or greater than those found in the population, on average, and that randomly assigning missing relatedness links is more likely to decrease heritability than non-randomly missing relatedness values, we conclude that background relatedness and missing pedigree links are unlikely to substantively decrease the heritability estimates obtained from our main models.

**S4. Glossary.**

For an in-depth study of back-transformation of parameters obtained through non-gaussian quantitative genetics GLMMs, we recommend Morrissey (2015); De Villemereuil *et al.* (2016); de Villemereuil (2018).

**Threshold model**. In quantitative genetics, a model where the phenotypic response variable is represented by ordered categories: a threshold model works by taking a categorical trait and creating a hypothetical "latent trait" (also called a “liability”) with an infinitesimal and gaussian distribution onto which the categorical trait is mapped. A single-threshold model applies to a phenotype with two categories, while a multiple-thresholds model has more than two categories that are ordered hierarchically (as is the case with ordinal data).

**Latent scale**. Scale at which a non-gaussian trait is hypothetically treated as a normally distributed (latent) trait (from De Villemereuil *et al.* (2016)). Applied to a quantitative genetic animal model, the distribution of the trait at this scale is a linear mixed model:

**l** = μ + **X**𝛃 + **Z_a_a** + **Z_1_u**_1_... + **Z_k_u_k_** + **o**	(S1)

where **l** is the vector of liabilities (phenotypes on the liability scale), μ is the model intercept, 𝛃 is the fixed effects vector, **a** are the additive genetic effects, **u_1_**, …, **u_k_**, are other random effects, while **X**, **Z_a_**, **Z_1_**,…, **Z_k_** are incidence matrices of the appropriate dimensions. The error **o** is normally distributed and is referred as “overdispersion".

**Expected scale**. For non-gaussian GLMMs, it is the scale at which every value on the latent scale is linked (by a link function) to an expected value on the scale of the observed trait. The formula linking the latent and the expected distribution of the trait depends on the link function. The trait is mapped onto the expected data scale by the inverse of the link function.

**Liability scale**. (Multi)Threshold models are an exception with respect to other non-gaussian GLMM: they technically do not have a latent scale, but a liability scale, since the threshold represents passage from one category of the trait to another. To derive heritability on the liability scale from the latent parameters in threshold models, it is necessary to add the "link variance" to the formula denominator when something different than “threshold” family and probit link is used. Otherwise, latent and liability estimates are identical, and liability values are deterministically mapped on the observed scale (there is no expected scale in (multi)threshold models). In other words, for threshold and ordinal models, estimates on the liability scale have a deterministic relationship with the trait (de Villemereuil 2012). As mentioned in the main text, the liability scale is currently more appropriate to report estimates obtained with animal models employing the ordinal family (de Villemereuil 2018).

**Observed/data/phenotypic scale**. Scale of the observed data. It is obtained from the expected scale by adding an error term around the expected value, to model the noise around it. Depending on the scale, the error term can have different distributions. This is the scale that represents the actual phenotype. Its relationship with the liability scale of the threshold model is different, as values on the liability scale are deterministically mapped on the observed data scale.

**Zero-inflated over-dispersed Poisson distribution**. A data distribution characterized by a high number of zeroes and a relatively lower number of finite count data. It can be analyzed via a generalized linear model that functions as a bivariate model, with the zero-inflation component (binary, logit scale) and a Poisson component (Poisson). For further details on the link formulas see (Bonnet *et al.* 2022).

**Lifetime breeding success**. A phenotypic trait detailing the number of progeny (zygotes) that an individual produced during the entire lifetime. For many long-lived species with high juvenile mortality, it can be analyzed through a zero-inflated Poisson model. Since it is often used to represent fitness, several studies use relative LBS (since relative fitness is a value used in evolutionary formulas). Note that while certain authors make a distinction between LBS and lifetime reproductive success, sensu McCleery *et al.* (2004), i.e., the number of individuals recruited as adults in the population, the two terms are often used synonymously. While in this study we use the term “Lifetime Breeding Success (LBS)” as it is used in Bonnet *et al.* (2022), note that Hendry *et al.* (2018) use the term “Lifetime Reproductive Success (LRS)” as a synonym of LBS, while using “Lifetime Recruitment Success (LrecS)” to indicate the number of individuals recruited in the adult population. In another instance, Van de Walle *et al.* (2022) used LBS for the number of offspring born, and LRS for the number of offspring weaned.

**V_A_**. Additive genetic variance.

**V_A_(*w*)**. Additive genetic variance in relative fitness. We estimated it following the methodology outlined in Bonnet *et al.* (2022), where the authors back-transformed the two liabilities obtained from the zero-inflation Poisson model in a single estimate of relative (not absolute/raw) fitness.

**Short-term fitness proxies**. Fitness proxies that span one generation (e.g., lifetime breeding success).

**Long-term fitness proxies**. Fitness proxies that span multiple generations (e.g., genetic contribution).

**Time-limited fitness proxies**. Short-term fitness proxies limited to a specific time window in the life of an individual (e.g., annual breeding success).

**S5: Variance-covariance random structure of bivariate models**

All six univariate models (annual breeding success, annual survival, annual fitness, triannual breeding success, LBS, and binary LBS) were fitted with an additive genetic effect. Thus, all bivariate models presented a variance-covariance matrix for the additive genetic effect:

$\mathbf{G}\otimes\mathbf{A}$	(S2)

$\mathbf{G}=\left| \begin{matrix} \text{σ}_{a_{D1}}^{\text{2}} & \text{σ}_{a_{D1D2}} \\ \text{σ}_{a_{D1D2}} & \text{σ}_{a_{D2}}^{\text{2}} \end{matrix} \right|$	(S3)

Where $\mathbf{G}$ is a 2x2 matrix representing additive genetic (co)variances for the pair of fitness metrics, and $\mathbf{A}$ is the additive genetic relationship matrix. Given fitness metric 1 and fitness metric 2, matrix $\mathbf{G}$ includes the variances $\text{σ}_{a_{D1}}^{\text{2}}$ (genetic variance of fitness metric 1), $\text{σ}_{a_{D2}}^{\text{2}}$(genetic variance of fitness metric 2), and the genetic covariance between metric 1 and 2, $\text{σ}_{a_{D1D2}}$. Note that while $\text{σ}_{a_{D1}}^{\text{2}}$and $\text{σ}_{a_{D2}}^{\text{2}}$ are also estimated in the univariate animal models of the corresponding fitness metrics, $\text{σ}_{a_{D1D2}}$ is unique to the bivariate analysis. In MCMCglmm variance-covariance matrices are fitted using the ‘us(trait):random_effect’ notation; an additive genetic variance-covariance matrix (such as Equation S2) would be written ‘us(trait):animal’, with “animal” being a reserved term of the function indicating the effect of the breeding value (genetic merit) of the individual. In addition to the additive genetic variance-covariance matrix, when bivariate models involved two fitness metrics that had other random effects in common (for example, univariate models investigating annual breeding success and annual survival were both fitted with a random effect of ‘hydroyear’), we estimated the covariances attributed to those random effects using the same matrix structure. In other words, our bivariate models also estimated the covariances between those random effects that were present in the univariate model of both fitness metrics. Consequently, any bivariate analysis where both fitness metrics included a permanent environmental effect also estimated the following variance-covariance matrix:

$\mathbf{Pe}\otimes\mathbf{I}$	(S4)

$\mathbf{Pe}=\left| \begin{matrix} \text{σ}_{\mathrm{Pe}_{D1}}^{\text{2}} & \text{σ}_{\mathrm{Pe}_{D1D2}} \\ \text{σ}_{\mathrm{Pe}_{D1D2}} & \text{σ}_{\mathrm{Pe}_{D2}}^{\text{2}} \end{matrix} \right|$	(S5)

with $\mathbf{Pe}$ as a 2x2 matrix representing permanent environmental (co)variances for the fitness metric pair and $\mathbf{I}$ as an identity matrix of appropriate size. The matrix $\mathbf{Pe}$ therefore includes the permanent environmental variances associated with fitness metrics 1 and 2, $\text{σ}_{\mathrm{Pe}_{D1}}^{\text{2}}$, and $\text{σ}_{\mathrm{Pe}_{D2}}^{\text{2}}$, and the covariance between the permanent environmental components of the fitness metrics, $\text{σ}_{\mathrm{Pe}_{D1D2}}$. In MCMCglmm this variance-covariance matrix can be fitted exactly like the additive genetic matrix, by using the ‘us(trait):random_effect’ formulation; the above matrix, for example, could be estimated by adding ‘us(trait):permanent_environment’ to the random effect structure of the model.

When a random effect was fitted only for one of the two fitness metrics, it was entered in the bivariate analysis with the ‘idh(at.level(trait,n)):random_effect’ formulation, where n is the position of the fitness metric that the random effects belongs to (either 1 or 2). For example, in the bivariate analysis of annual survival and triannual breeding success, since the random effect of hydroyear is present only in the model for annual survival (fitness metric 1), and not in the model for triannual breeding success (fitness metric 2), the random effect structure was: ‘idh(at.level(trait,1)):hydroyear’, and the resulting matrix representing hydroyear variance-covariances ($\mathbf{HY}$) would be a simple:

$\mathbf{HY}=\left| \begin{matrix} \text{σ}_{\mathrm{HY}_{D1}}^{\text{2}} & 0 \\ \text{0} & \text{0} \end{matrix} \right|$	(S6)

**S6: Full list of random effects structures of bivariate models**

We iterated all bivariate models that did not involve LBS and binary LBS a sufficient number of times to obtain minimum effective sample sizes from the posterior of 6114 for random effects and 7602 for fixed effects. For the bivariate animal models involving pairs of annual fitness metrics (annual survival x annual breeding success; annual survival x annual fitness; annual fitness x annual breeding success) the random effects structure was:

us(trait):animal +

us(trait):permanent_environment +

us(trait):hydroyear +

us(trait):hydroyear_group +

us(trait):mother_id +

us(trait):units, where the last term refers to the residual variance.

For bivariate animal models involving pairs composed by an annual fitness metric and triannual breeding success (annual survival x triannual breeding success; annual breeding success x triannual breeding success; annual fitness x triannual breeding success) the random effects structure was:

us(trait):animal +

us(trait):permanent_environment +

idh(at.level(trait,1)):hydroyear +

idh(at.level(trait,1)):hydroyear_group +

us(trait):mother_id +

us(trait):units.

For bivariate animal models involving pairs composed by an annual fitness metric and binary LBS (annual survival x binary LBS; annual breeding success x binary LBS; annual fitness x binary LBS) the random effects structure was:

us(trait):animal +

idh(at.level(trait,1)):permanent_environment +

idh(at.level(trait,1)):hydroyear +

idh(at.level(trait,1)):hydroyear_group +

idh(at.level(trait,2)):cohort +

us(trait):mother_id +

us(trait):units.

For the bivariate animal model involving the pair composed by triannual breeding success x binary LBS the random effects structure was:

us(trait):animal +

us(at.level(trait,1)):permanent_environment +

us(at.level(trait,2)):cohort +

us(trait):mother_id +

us(trait):units.

For bivariate models involving LBS (which are technically trivariate models, because zero-inflated Poisson models have two components) we took several steps to increase effective sample size, decrease processing times, and help convergence. Specifically, we excluded all fixed and random parameters that never influenced results in other models, and we parallelized each run between 10 chains (Wolak & Reid 2017). We obtained minimum effective sample sizes of 1114 for random effects and 1691 for fixed effects. For bivariate animal models involving pairs composed by an annual fitness metric and LBS (annual survival x LBS; annual breeding success x LBS; annual fitness x LBS) the random effects structure was simplified with respect to the full random structure by eliminating random effects that never showed non-zero variances:

us(trait):animal +

idh(at.level(trait,1)):hydroyear +

idh(at.level(trait,1)):hydroyear_group +

us(trait):units.

For the bivariate animal model involving the pair composed by triannual breeding success x LBS the random effects structure was simplified with respect to the full random structure by eliminating random effects that never showed non-zero variances:

us(trait):animal +

us(trait):units.

For the bivariate animal model involving the pair composed by LBS x binary LBS the random effects structure was:

us(trait):animal +

us(trait):cohort +

us(trait):mother_id +

us(trait):units.

**S7: Permutation of bivariate analyses**

To further test the capability of our bivariate models to recover accurate genetic correlation estimates we ran the annual breeding success x annual survival bivariate analysis with a permuted dataset. The annual breeding success and annual survival records of all individuals were kept the same, but the positions of the individuals in the pedigree were permuted. All other parameters of the model were kept exactly as described in the main text, except for the number of iterations in the model and in the back-transformation, which were reduced to save processing time. We obtained a much lower genetic correlation than with the non-permuted pedigree (genetic correlation (posterior modes [credible intervals]) in the original analysis r_a_ = 0.9866 [0.8228, 0.9974]; with permuted pedigree: r_a_ = 0.0257 [-0.9622, 0.7858]). As expected given our permutation scheme, the phenotypic correlation did not change nearly as dramatically (original analysis: r_p_ = 0.0450 [0.0187, 0.0599]; with permuted pedigree: r_p_ = 0.0341 [0.0068, 0.0543]). The reason for this difference is that phenotypic correlations are not influenced by the genetic relationships between individuals, while genetic correlations are. To test another type of permutation on a different bivariate model, we also ran a bivariate analysis between annual breeding success and a permuted binary LBS: all individuals this time had their correct annual breeding success records and the correct position in the pedigree, but the binary LBS record was permuted. Again, we obtained posterior estimates for the genetic correlation that substantially overlapped 0 (original analysis: r_a_ = 0.8908 [0.2090, 0.9988]; permuted binary LBS: r_a_ = 0.1514 [-0.7240, 0.8108]).

SUPPLEMENTARY RESULTS

**S8. V_A_(*w*) estimates obtained in Amboseli baboons**

There are differences of opinion concerning how to best report the additive genetic component of fitness (Hendry *et al.* 2018; Bonnet *et al.* 2019a). In our study we decided to report both additive genetic variance and heritability of (absolute) lifetime fitness (LBS, or absolute LBS), to facilitate generalizations and comparisons with other studies and with time-limited fitness metrics. However, when discussing lifetime fitness metrics, reporting the genetic variance of relative lifetime fitness (V_A_(*w*)) can also be informative (Bonnet *et al.* 2019a).

Bonnet *et al.* (2022) estimated V_A_(*w*) for several long-term study populations, including the baboons that are the subjects of our study. We followed the protocol they developed to also estimate V_A_(*w*) with our LBS univariate model, with the goal of comparing between our estimate of V_A_(*w*) and the one obtained in their study. However, the ultimate aim of the Bonnet *et al.* (2022) study was to obtain a meta-analytic average of V_A_(*w*), using both sexes in each population. For this reason, the authors chose fixed and random effects that could be applied across several different study systems. Since our focus was solely on female baboons, we chose parameters that were population-specific. In addition to including the fixed and random effects described in the main text, our analysis of LBS differed from that of Bonnet *et al.* (2022) because we did not include the fixed effects of social dominance, inbreeding, and genetic groups (a factor that models the expected proportion of immigrant genetic ancestry to control for unknown parents belonging to potentially genetically differentiated populations, sensu Wolak and Reid (2017); Bonnet *et al.* (2022)). Social dominance is known to affects fitness in several primates (Silk 2007; Blomquist *et al.* 2011; Snyder-Mackler *et al.* 2020), even though in baboons the relationship is not straightforward (Campos *et al.* 2020). However, we did not include this effect as we have evidence that agonistic behaviour, which determines social dominance in our population, has a sizable additive genetic component (McLean et al., in prep). While extricating the social and genetic components of inheritance-based rank is a challenge, we suspect that social dominance might genetically covary with fitness; in that case, it would be more appropriate to investigate the role of rank on fitness using a full bivariate analysis (Wilson 2008). From the lifetime fitness models we also excluded covariates representing inbreeding, as inbreeding estimated from the pedigree has been shown to be very low in this population (Bonnet *et al.* 2022); and a covariate representing genetic groups, as our population does not have a clear generation zero “base population”, no females immigrated from other populations, and many unknown fathers originate from the same ecosystem, and are unknown simply because they were born in groups that are not intensively monitored. In addition, neither covariate influenced lifetime fitness in the analyses by Bonnet *et al.* (2022).

Bonnet *et al.* (2022) also included the random interaction of mother:cohort. We chose not to do so here because in baboons, mothers have only one surviving infant per year and thus this random effect has inevitably as many levels as the number of female baboons in the dataset. We believe that this specific difference is crucial to the comparison of our results, as the additive genetic variance of relative fitness that we obtained (V_A_(*w*) = 0.902 [0.491, 1.364]) was higher than that obtained by Bonnet et al. (V_A_(*w*) = 0.231 [0.0314, 0.547]), but they also obtained a sizable variance for the mother:cohort interaction (V_M:C_(*w*) = 0.285 [0.128, 0.664]. Repeating their analysis (with the dataset they provided) without the random effect of mother:cohort nearly doubled the estimate to a V_A_(*w*) = 0.445, still lower than ours but more comparable.

The other main reason for the difference between our estimates of V_A_(*w*) is, of course, the presence of males in the dataset used by Bonnet *et al.* (2022). The V_A_(*w*) value we found in female refers to only half the population, and if we were to consider LBS a uniquely female trait, the actual evolutionary response would be halved. While data on male LBS are incomplete, the offspring numbers for males that are currently available and that were used by Bonnet *et al.* (2022) can provide an unbiased, if imprecise, estimate of male LBS. The lower V_A_(*w*) obtained when males were included is compatible with several possibilities. The estimates of male LBS may be too imperfect to estimate V_A_(*w*) well; alternatively, there could be antagonistic pleiotropy for LBS in the Amboseli baboons. In addition, if the V_A_(*w*) of male baboons was lower than that of females, this would likely be enough to decrease the overall V_A_(*w*), even without sexually antagonistic pleiotropy. Finally, a non-mutually exclusive explanation is that the social factors that could lead to inflated estimates of LBS heritability in females might not affect males in the same way: by leaving the natal group at sexual maturity, the influence of maternal kin is reduced for males compared to females.

**S9. Expected increase of relative fitness in Amboseli baboons**

V_A_(*w*) can also be used to estimate the maximum rate of expected increase in LBS in our population during our study period, i.e., the expected growth in mean LBS that would have happened if natural selection translated all genetic variance in fitness into phenotypic change. Although our study population has slightly grown since the 1970s – possibly recovering after a drastic reduction in the 1960s (Altmann *et al.* 1985) – this increase has been several order of magnitudes below what our genetic parameters indicate. The formula detailed in Bonnet et al. (2022, Supplementary Text S3 “Relative speed of adaptation”) used to calculate $\overline{w}$, the ratio between the average relative fitness in the base population and the average relative fitness after *g* generations, is:

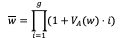
 (S7)

Entering the V_A_(*w*) = 0.902 estimated by our analysis of LBS, we calculated that if the genetic potential had translated into a corresponding phenotypic change, after only two generations our mean relative LBS should have increased of a factor of 5.32, i.e. the average LBS of the granddaughters of females born in the 1970 should have been more than five times higher (note that we assume here an equal V_A_(*w*) for the entire population, not only females). The increase in average fitness was instead small relative to that predicted by the genetic potential (from 1.636 average LBS of females born in the 1970’s to 2.094 average LBS for females born in the 1990’s), which confirms that, in females of our population, phenotypic growth in mean LBS does not correspond to the genetic evolutionary potential (Bonnet *et al.* 2022).

Overall, we note that the importance of Bonnet *et al.* (2022) study does not lie only in the estimates they report, but also in the detailed record of the methodology and in the public availability of code and datasets, without which this comparison could not have been made. The authors’ efforts to share all stages of their work created opportunities for further, in-depth analyses and comparisons such as the one we reported, which expand on their results and advance the field in a way that would not otherwise be possible.

**S10. Non-genetic sources of variance and fixed effects**

All time-limited fitness metrics increased with linear age (or triennium) but decreased in later life (i.e., all showed a quadratic relationship with age or triennium). Annual survival increased over the first 4 years of life of female baboons, and then started decreasing again around age 12 (Figure S1); annual breeding success rapidly increased from age 5 to age 9 and then decreased consistently after age 17 (Figure S2). High rainfall in the hydroyear preceding the target hydroyear was associated with lower annual breeding success and lower annual fitness, i.e., females had lower annual breeding success following hydroyears with higher rainfall and higher annual breeding success following hydroyears with lower rainfall. We speculate that decreased breeding success during drought hydroyears might have made females more likely to produce an offspring in the hydroyears immediately following (Lea *et al.* 2015). Conversely, hydroyears with average or high rainfall might have promoted reproduction, which would then have decreased the chances of a second birth in the following hydroyears. An alternative approach to modeling rainfall in the year prior to the target year would be to model rainfall in the current hydroyear. However, when we explored this alternative specification, a fixed effect of rainfall did not significantly predict any fitness metric. Better-quality habitat was associated with increased breeding success and annual fitness, but surprisingly also with decreased annual survival, even though our dataset included a higher relative proportion of female-years where death occurred in the worse-quality habitat. The reason for this result was probably that the two groups transferred to the better-quality habitat after living in the worse-quality one. Thus, many individuals who were born in the worse-quality habitat died in the better-quality one, whereas it was impossible for our study subjects to be born in the better-quality habitat and die in the worse one (Supplementary Table S3). Group size did not have any effect on time-limited fitness metrics, and as expected, no fixed predictor had any effect on lifetime fitness metrics (Supplementary Table S3).

The random effects of hydroyear explained non-zero variance for annual breeding success and annual fitness, but not for annual survival (Table 2, Figure 2). Reproduction and the first year of life of an infant are a particularly sensitive time, which might make annual breeding success more sensitive to environmental conditions than annual survival. However, we also found group-hydroyear effects for all three annual fitness metrics (Table 2, Figure 2), which suggests that group-level processes can influence both female baboons’ survival and breeding success. Resources in Amboseli vary not only over time but also over space, and while groups often occupy territories that geographically overlap, a finer, temporally differentiated partition of the landscape might still be linked to group differences in resource availability (Markham *et al.* 2013). For example, two groups might share the same territory, but within it, one group might have clear priority of access to an ephemeral waterhole or to the best foraging grounds. While we can only speculate what mechanisms might cause hydroyear and group-hydroyear variations in fitness, we note that the greatest portion of phenotypic variance was by far the residual variance (i.e., stochastic error), which indicates that most of the forces acting on fitness in this population are yet to be quantified.

**S11: Heritability of fitness metrics estimated from bivariate models**

The bivariate models estimated heritability values similar overall to those recovered via univariate models, such that the credible intervals of most of corresponding heritability estimates overlapped with each other. An exception was the heritability of annual breeding success in the bivariate models: despite being very low in all models, it often differed from zero in bivariate but not univariate models, with estimates of the posterior modes ranging from h^2^_observed_ = 0.0011 to h^2^_observed_ = 0.0223 in the bivariate models (Table S4). Heritability of annual survival also varied among the bivariate models, ranging from h^2^_observed_ = 0.1120 to h^2^_observed_ = 0.3839; notably, the average value across the bivariate models (mean h^2^_observed_ = 0.2327) was very similar to the estimate recovered from the univariate analysis (h^2^_observed_ = 0.2317). Heritability estimates obtained for binary LBS and LBS had credible intervals that overlapped substantially with those obtained by the univariate models (Table 3 and S4). Estimates for fixed effects and non-genetic random effects were largely consistent between bivariate and univariate analyses.

**S12: Evolutionary change in annual survival**

To assess evolutionary change in annual survival we tested if the rate of change in the estimated breeding values (EBVs, the estimated genetic “merit” of an individual for a trait) differed from what could be expected under a null model of evolutionary change that included only drift. To do so, we followed the protocol detailed in Hadfield *et al.* (2010) and Bonnet *et al.* (2019b), which is widely used (Bonnet *et al.* 2017; Tuliozi *et al.* 2023) and was found to be consistent with genomic metrics of rate of change (Hunter *et al.* 2022). First, we extracted the EBVs for all individuals from the univariate model of annual survival. Second, we regressed the EBVs against hydroyear of birth to estimate the rate of change in the EBVs in Amboseli baboon females over time. We repeated this process across all iterations of the model, obtaining a vector of regression coefficients (slopes) representing re-sampled rates of change in the EBVs of our population. Then we compared this slope distribution with a vector of corresponding length containing the rates of change in the EBVs of our population when only evolutionary change by drift was considered. In brief, the drift-only model uses the genetic variance associated with the trait (in this case annual survival) to drop simulated EBVs down the pedigree, using the same general assumptions of the animal model. The posterior mode of the rate of change in the EBVs for annual survival was 0.013 [0.006, 0.022]. 99.05% of the replicates had a more positive slope than expected under the null model of evolutionary change by drift alone (Supplementary Figure S3). This constitutes strong evidence for evolutionary genetic change in annual survival (Bonnet *et al.* 2017). On the data scale, this change can be translated to an average expected increase of 14.8% in annual survival probability between individuals born in 1971 and those born in 2022 (Supplementary Figure S4).

Populations at equilibrium are predicted to have very low V_A_ for fitness and fitness-correlated traits, as purifying selection is expected to deplete their genetic variance. However, changes in environmental conditions can maintain genetic variance through fluctuating selective pressures, even across timespans as short as two generations (Bergland *et al.* 2014; Bitter *et al.* 2024). A history of fluctuating selection is possible for our study population, given the very dynamic nature of the Amboseli ecosystem (Western & Van Praet 1973; Western & Behrensmeyer 2009; Alberts & Altmann 2012), and it might partially explain why our study population presents substantial genetic variation for crucial traits like annual survival. However, note that the high V_A_ for fitness and the positive change in the EBVs for annual survival during our study period suggest a successful response to directional selection rather than frequent environment-dependent changes in the sign and magnitude of the relationship between traits and fitness. We favor this interpretation because genetic diversity maintained by fluctuating selection implies fluctuating change(s) in the correlation between trait(s) and fitness, but when the trait examined is fitness itself, finding V_A_ for fitness implies that the relationship between genotype and fitness did not change over time. In other words, environmental fluctuation can maintain overall genetic variance, but it also tends to depress estimates of V_A_ in fitness when fitness is averaged across fluctuations. We note, however, that while our estimates suggest that directional selection has been consistent during the study period, environmental changes before our study began could have modified the relationship between crucial traits and fitness, preserving genetic variance in the traits that give rise to differences in fitness. Thus, while our current time window captures a phase of directional selection, a legacy of fluctuating selection could help explain the substantial genetic variation we observe today.

SUPPLEMENTARY TABLES

**Table S1**. **Univariate animal models including genetic ancestry scores: posterior modes and 95% credible intervals of all fixed effects that influence fitness metrics in univariate animal models (QGGLMMs), using the sub-sample of the dataset for which genetic ancestry scores were available**. The effects are reported on the latent scale and are marked in **bold** when the 95% credible intervals do not contain 0.

| Trait | Variable | Posterior mode | 95% CI |
| --- | --- | --- | --- |
| Annual breeding success | **(Intercept)** | **-1.3792** | **[-1.6746, -1.1519]** |
|  | Genetic ancestry (z-transf) | -0.0299 | [-0.1008, 0.0391] |
|  | **Age (z-transf)** | **3.1015** | [2.7420, 3.4431] |
|  | **Age^2^ (z-transf)** | **-2.3638** | **[-2.6286, -2.0523]** |
|  | Group size (z-transf) | 0.0707 | [-0.2744, 0.4368] |
|  | (Group size)^2^ (z-transf) | -0.0595 | [-0.4617, 0.2470] |
|  | **Rainfall anomaly in previous year (z-transf)** | **-0.1489** | **[-0.2288, -0.0489]** |
|  | Habitat quality | -0.0058 | [-0.2759, 0.2647] |
| Annual survival | **(Intercept)** | **5.1491** | **[3.3378, 8.3916]** |
|  | Genetic ancestry (z-transf) | 0.0732 | [-0.4777, 0.5931] |
|  | Age (z-transf) | -0.5451 | [-1.6560, 0.2584] |
|  | **Age^2^ (z-transf)** | **-1.0485** | **[-1.8479, -0.3563]** |
|  | Group size (z-transf) | -0.2245 | [-1.5615, 1.2693] |
|  | (Group size)^2^ (z-transf) | 0.3804 | [-0.9838, 1.9725] |
|  | Rainfall anomaly in previous year (z-transf) | -0.2088 | [-0.4496, 0.0605] |
|  | **Habitat quality** | **-1.4680** | **[-3.4881, 0.0800]** |
| Annual fitness | **(Intercept)** | **2.7018** | **[2.3371, 2.9858]** |
|  | Genetic ancestry (z-transf) | -0.0085 | [-0.0953, 0.0714] |
|  | **Age (z-transf)** | **2.4286** | **[2.2113, 2.7465]** |
|  | **Age^2^ (z-transf)** | **-2.0745** | **[-2.3175, -1.8468]** |
|  | Group size (z-transf) | 0.1100 | [-0.3334, 0.5290] |
|  | (Group size)^2^ (z-transf) | -0.0402 | [-0.5318, 0.3428] |
|  | **Rainfall anomaly in previous year (z-transf)** | **-0.1658** | **[-0.3124, -0.0586]** |
|  | Habitat quality | -0.0067 | [-0.3813, 0.2784] |
| Triannual breeding success | **(Intercept)** | -0.3572 | [-0.8241, 0.0850] |
|  | Genetic ancestry (z-transf) | -0.0532 | [-0.1950, 0.1280] |
|  | **Triennium (z-transf)** | **6.7616** | **[5.8935, 7.3833]** |
|  | **Triennium^2^ (z-transf)** | **-5.5042** | **[-6.1266, -4.8060]** |
|  | Group size (z-transf) | 0.2593 | [-0.6116, 0.9136] |
|  | (Group size)^2^ (z-transf) | -0.2627 | [-0.9386, 0.5466] |
|  | Previous rainfall anomaly (z-transf) | -0.1043 | [-0.2452, 0.0406] |
|  | **Habitat quality** | **-0.5758** | **[-1.0015, -0.0555]** |
| Binary LBS | **(Intercept)** | **3.5311** | **[1.0806, 7.5516]** |
|  | Genetic ancestry (z-transf) | 0.7561 | [-0.7821, 3.7813] |
|  | Rainfall anomaly in year before birth (z-transf) | 0.2352 | [-0.8772, 1.4590] |
|  | Habitat quality in year before birth | 1.3680 | [-3.7509, 7.7463] |
|  | Cohort (z-transf) | -0.6871 | [-5.3671, 2.0418] |
| LBS, Poisson | **(Intercept Poisson)** | **1.5379** | **[1.0667, 1.8862]** |
|  | Genetic ancestry (z-transf) | 0.0535 | [-0.1237, 0.2329] |
|  | Rainfall anomaly in year before birth (z-transf) | 0.0190 | [-0.1565, 0.1977] |
|  | Habitat quality in year before birth | 0.0703 | [-0.7728, 0.6889] |
|  | Cohort (z-transf) | -0.1450 | [-0.5273, 0.3000] |
| LBS, zero-inflated | **(Intercept zero-inflated)** | **-19.6089** | **[-20.1912, -18.9290]** |
|  | Genetic ancestry (z-transf) | 0.0273 | [-0.2910, 0.3359] |
|  | Rainfall anomaly in year before birth (z-transf) | 0.0081 | [-0.2560, 0.2766] |
|  | Habitat quality in year before birth | -0.4327 | [-1.4184, 0.8980] |
|  | Cohort (z-transf) | 0.1353 | [-0.5332, 0.8260] |

**Table S2**. **Model parameters for both univariate and bivariate animal models**. Thinning, number of iterations, and length of burn-in all refer to the MCMC iterations stored by the MCMCglmm for both univariate (Fitness trait) and bivariate (Fitness trait 1 x Fitness trait 2) models.

| Model | Thinning | Number of iterations | Length of burn-in |
| --- | --- | --- | --- |
| Annual breeding success | 500 | 5500000 | 500000 |
| Annual survival | 1000 | 11000000 | 1000000 |
| Annual fitness | 500 | 5500000 | 500000 |
| Triannual breeding success | 500 | 5500000 | 500000 |
| Binary LBS | 500 | 5500000 | 500000 |
| LBS | 1000 | 16000000 | 1000000 |
| Annual breeding success x Annual survival | 500 | 5500000 | 500000 |
| Annual breeding success x Annual fitness | 500 | 5500000 | 500000 |
| Annual breeding succ. x Triannual breeding succ. | 500 | 5500000 | 500000 |
| Annual breeding success x Binary LBS | 750 | 11000000 | 1000000 |
| Annual breeding success x LBS† | 4000 | 51000000 | 1000000 |
| Annual survival x Annual fitness | 500 | 5500000 | 500000 |
| Annual survival x Triannual breeding success | 500 | 5500000 | 500000 |
| Annual survival x Binary LBS | 750 | 11000000 | 1000000 |
| Annual survival x LBS^a^ | 4000 | 51000000 | 1000000 |
| Annual fitness x Triannual breeding success | 500 | 5500000 | 500000 |
| Annual fitness x Binary LBS | 750 | 11000000 | 1000000 |
| Annual fitness x LBS^a^ | 4000 | 51000000 | 1000000 |
| Triannual breeding success x Binary LBS | 750 | 11000000 | 1000000 |
| Triannual breeding success x LBS^a^ | 4000 | 51000000 | 1000000 |
| Binary LBS x LBS | 4000 | 6000000 | 1000000 |

^a^: models with LBS are composed of ten parallelized chains, each with 6000000 iterations, thinning of 4000 and burn-in of 1000000.

**Table S3**. **Univariate animal models:** **posterior modes and 95% credible intervals for all fixed effects included in models of different fitness metrics, estimated via univariate animal models (QGGLMMs)**. The effects are reported on the latent scale and are marked in **bold** when the 95% credible intervals do not contain 0.

| Trait | Variable | Posterior mode | 95% CI |
| --- | --- | --- | --- |
| Annual breeding success | **(Intercept)** | **-1.7081** | **[-1.9083, -1.5386]** |
|  | **Age (z-transf)** | **3.3021** | **[3.0736, 3.5994]** |
|  | **Age^2^ (z-transf)** | **-2.5479** | **[-2.7789, -2.3243]** |
|  | Group size (z-transf) | 0.0380 | [-0.3236, 0.3010] |
|  | (Group size)^2^ (z-transf) | -0.0666 | [-0.3634, 0.2764] |
|  | **Rainfall anomaly in previous year (z-transf)** | **-0.1476** | **[-0.2249, -0.0668]** |
|  | **Habitat quality** | **0.2459** | **[0.0639, 0.4493]** |
| Annual survival | (Intercept) | 1.0288 | [0.6817, 1.3847] |
|  | **Age (z-transf)** | **0.6151** | **[0.3540, 0.8552]** |
|  | **Age^2^ (z-transf)** | **-0.9177** | **[-1.1114, -0.7255]** |
|  | Group size (z-transf) | 0.1749 | [-0.4022, 0.6341] |
|  | (Group size)^2^ (z-transf) | -0.0937 | [-0.6892, 0.3971] |
|  | Rainfall anomaly in previous year (z-transf) | -0.0351 | [-0.1423, 0.0618] |
|  | **Habitat quality** | **-0.3527** | **[-0.7731, -0.0350]** |
| Annual fitness | (Intercept) | 1.9685 | [1.7443, 2.1680] |
|  | **Age (z-transf)** | **2.5423** | **[2.3485, 2.6853]** |
|  | **Age^2^ (z-transf)** | **-2.1111** | **[-2.2778, -1.9469]** |
|  | Group size (z-transf) | 0.0552 | [-0.2834, 0.4380] |
|  | (Group size)^2^ (z-transf) | -0.1168 | [-0.4815, 0.2695] |
|  | **Rainfall anomaly in previous year (z-transf)** | **-0.1416** | **[-0.2284, -0.0339]** |
|  | **Habitat quality** | **0.2642** | **[0.0348, 0.4966]** |
| Triannual breeding success | (Intercept) | -1.2224 | [-1.4865, -0.9004] |
|  | **Triennium (z-transf)** | **6.9667** | **[6.4977, 7.6138]** |
|  | **Triennium^2^ (z-transf)** | **-5.9073** | **[-6.3836, -5.3568]** |
|  | Group size (z-transf) | -0.3144 | [-0.9300, 0.2269] |
|  | (Group size)^2^ (z-transf) | 0.2019 | [-0.3374, 0.8185] |
|  | Previous rainfall anomaly (z-transf) | -0.0740 | [-0.1713, 0.0242] |
|  | Habitat quality | 0.0292 | [-0.2684, 0.3185] |
| Binary LBS | **(Intercept)** | **-1.4088** | **[-2.8243, -0.2690]** |
|  | Rainfall anomaly in year before birth (z-transf) | 0.0172 | [-0.5241, 0.4313] |
|  | Habitat quality in year before birth | 0.2070 | [-1.4176, 1.7962] |
|  | Cohort (z-transf) | -0.5527 | [-1.5667, 0.1774] |
| LBS, Poisson | **(Intercept Poisson component)** | **1.4174** | **[0.9244, 1.7902]** |
|  | Rainfall anomaly in year before birth (z-transf) | -0.0199 | [-0.1820, 0.1292] |
|  | Habitat quality in year before birth | -0.1920 | [-0.7581, 0.4067] |
|  | Cohort (z-transf) | 0.0724 | [-0.1902, 0.4037] |
| LBS, zero-inflated | **(Intercept zero-inflated component)** | **2.7607** | **[0.3294, 5.2755]** |
|  | Rainfall anomaly in year before birth (z-transf) | -0.0446 | [-1.1621, 1.2761] |
|  | Habitat quality in year before birth | 0.1014 | [-3.1173, 3.6204] |
|  | Cohort (z-transf) | 1.2459 | [-0.4490, 3.3665] |

**Table S4**. **Bivariate animal models: posterior modes and 95% credible intervals for all observed heritability values obtained from bivariate models of different fitness metrics**. Trait 1 and Trait 2 refer to the two traits in the bivariate model. h^2^ on the observed scale refers to Trait 1 and is marked in **bold** when the lower bound of the 95% CI > 0.001. Values below 10^-4^ are marked 0.

| Trait 1 Trait 2 | | h^2^ | 95% CI |
| --- | --- | --- | --- |
| **Annual breeding success** | (Annual survival) | **0.0067** | **[0.0022, 0.0122]** |
| **Annual** **breeding success** | (Annual fitness) | **0.0091** | **[0.0052, 0.0144]** |
| **Annual** **breeding success** | (Triannual breeding success) | **0.0061** | **[0.0023, 0.0105]** |
| Annual breeding success | (Binary LBS) | 0.0011 | [0, 0.0054] |
| **Annual breeding success** | (LBS) | **0.0223** | **[0.0074, 0.0410]** |
| **Annual survival** | (Annual breeding success) | **0.1910** | **[0.1111, 0.2573]** |
| **Annual survival** | (Annual fitness) | **0.3839** | **[0.3239, 0.4157]** |
| **Annual survival** | (Triannual breeding success) | **0.2094** | **[0.1383, 0.2764]** |
| **Annual survival** | (Binary LBS) | **0.2671** | **[0.2193, 0.3068]** |
| **Annual survival** | (LBS) | **0.1120** | **[0.0360, 0.1352]** |
| **Annual fitness** | (Annual breeding success) | **0.0125** | **[0.0077, 0.0144]** |
| **Annual fitness** | (Annual survival) | **0.1358** | **[0.1017, 0.1703]** |
| **Annual fitness** | (Triannual breeding success) | **0.0201** | **[0.0130, 0.0295]** |
| **Annual fitness** | (Binary LBS) | **0.0206** | **[0.0102, 0.0291]** |
| **Annual fitness** | (LBS) | **0.0746** | **[0.0521, 0.1423]** |
| **Triannual breeding success** | (Annual breeding success) | **0.0201** | **[0.0058, 0.0313]** |
| **Triannual breeding success** | (Annual survival) | **0.0280** | **[0.0154, 0.0450]** |
| **Triannual breeding success** | (Annual fitness) | **0.0728** | **[0.0536, 0.0968]** |
| **Triannual breeding success** | (Binary LBS) | **0.0168** | **[0.0028, 0.0250]** |
| **Triannual breeding success** | (LBS) | **0.1141** | **[0.1989, 0.3019]** |
| **Binary LBS** | (Annual breeding success) | **0.3758** | **[0.2528, 0.4535]** |
| **Binary LBS** | (Annual survival) | **0.4414** | **[0.3837, 0.4916]** |
| **Binary LBS** | (Annual fitness) | **0.3988** | **[0.3121, 0.4510]** |
| **Binary LBS** | (Triannual breeding success) | **0.3667** | **[0.2527, 0.4507]** |
| **Binary LBS** | (LBS) | **0.4496** | **[0.3809, 0.4776]** |
| **LBS** | (Annual breeding success) | **0.2502** | **[0.2111, 0.3139]** |
| **LBS** | (Annual survival) | **0.2968** | **[0.2563, 0.3498]** |
| **LBS** | (Annual fitness) | **0.2586** | **[0.2232, 0.3213]** |
| **LBS** | (Triannual breeding success) | **0.2419** | **[0.1989, 0.3019]** |
| **LBS** | (Binary LBS) | **0.3566** | **[0.2736, 0.4776]** |

**Supplementary Figures**

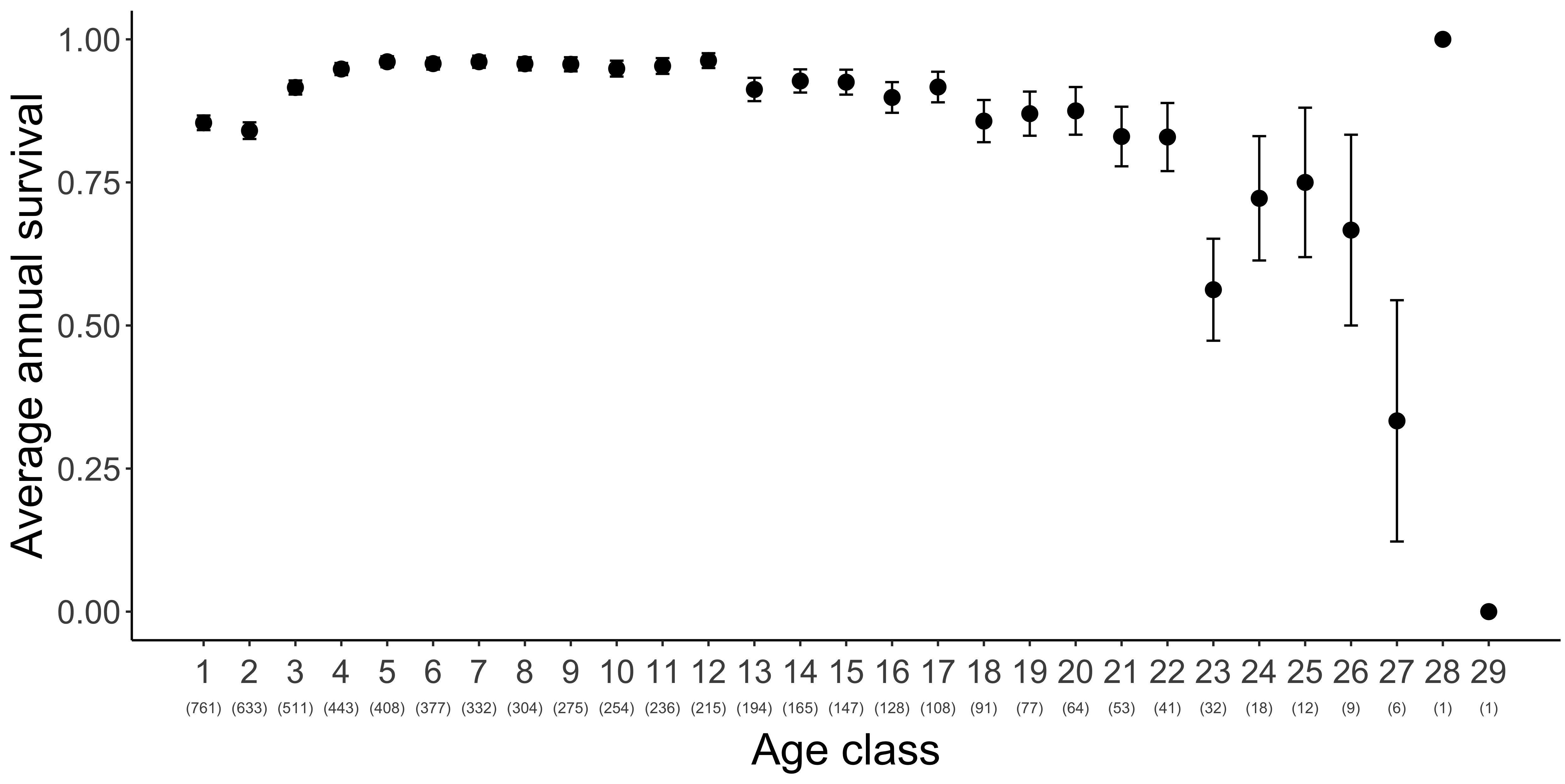

**Figure S1**. **Average annual survival as a function of age class in 806 female baboons**. Annual survival is recorded during each hydroyear (1^st^ Nov – 31^st^ Oct) of a female’s life: we show hydroyearly means and standard errors. Sample sizes for each age class are reported in parentheses below the corresponding age class. Note that the higher survival in the first age class relative to the second reflects the fact that the first female-hydroyear is shorter than the second for most females in the dataset (unless they were born on the 1^st^ of November, their first year did not encompass a full calendar year).

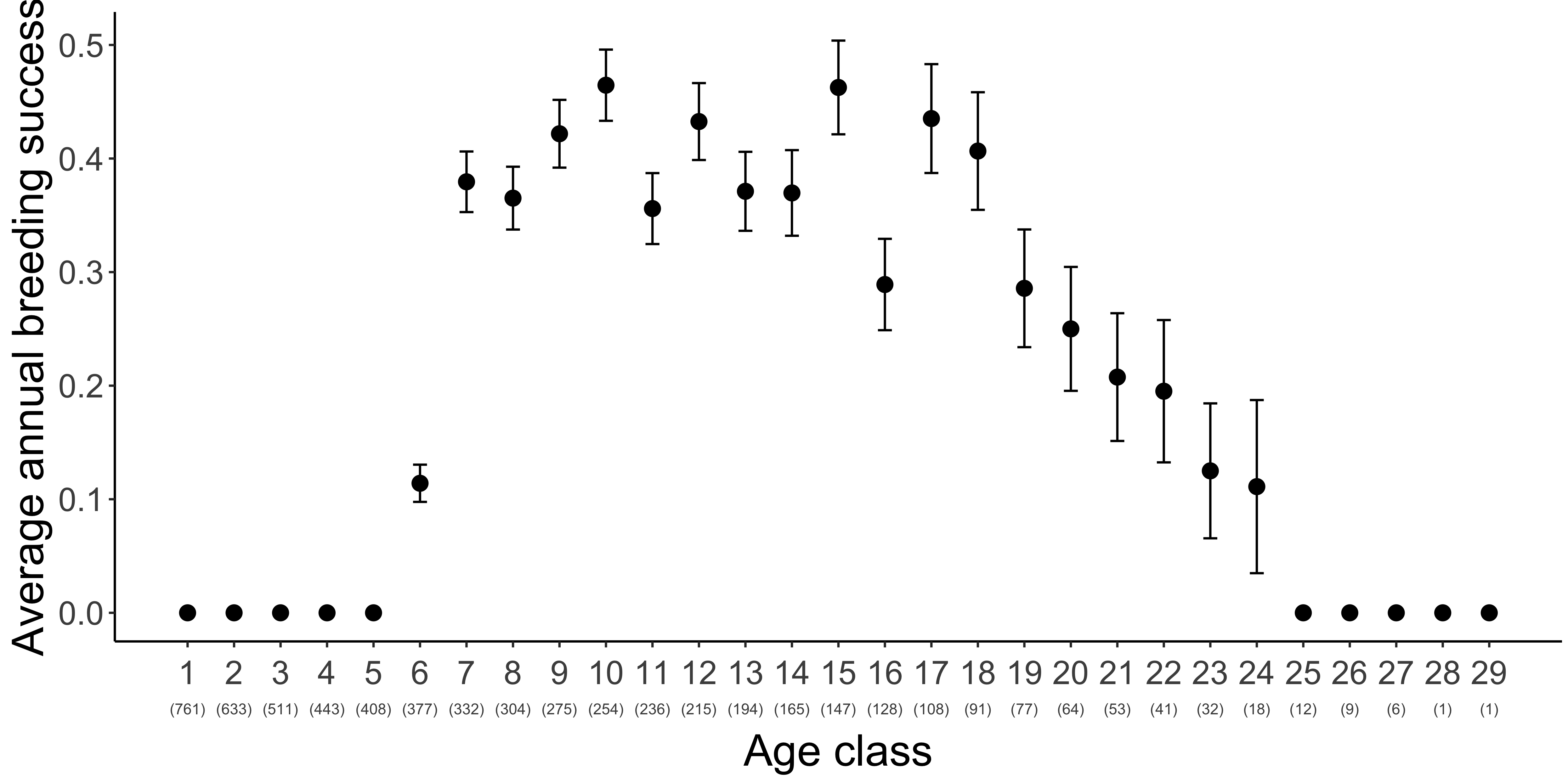

**Figure S2**. **Average annual breeding success as a function of age class in 806 female baboons**. Annual breeding success is recorded during each hydroyear (1^st^ Nov – 31^st^ Oct) of a female’s life: we show hydroyearly means and standard errors. Sample sizes for each age class are reported in parentheses below the corresponding age class.

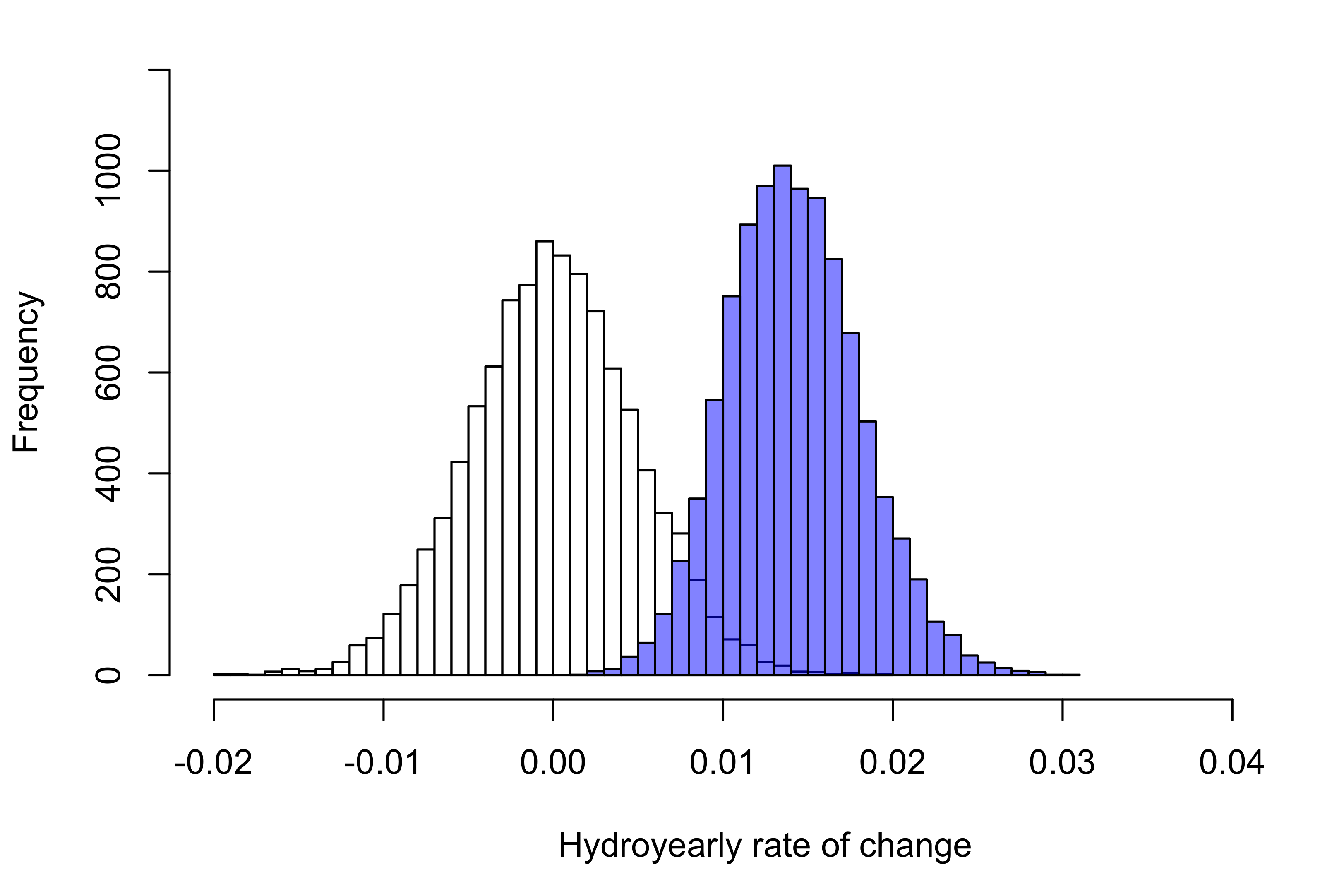

**Figure S3**. **Distribution of regression coefficients (slopes) for change in estimated breeding values (EBVs) of annual survival.** For each MCMC replicate, the rate of change (slope) in the EBVs of annual survival over time was calculated by regressing the average cohort EBVs (i.e., the EBVs of individuals born in the same hydroyear) over the hydroyear. These slopes represent the average change in the EBVs of the population happening over a hydroyear and are plotted with blue bars. Slopes obtained from a null model of evolutionary change that considers drift only are plotted with white bars.

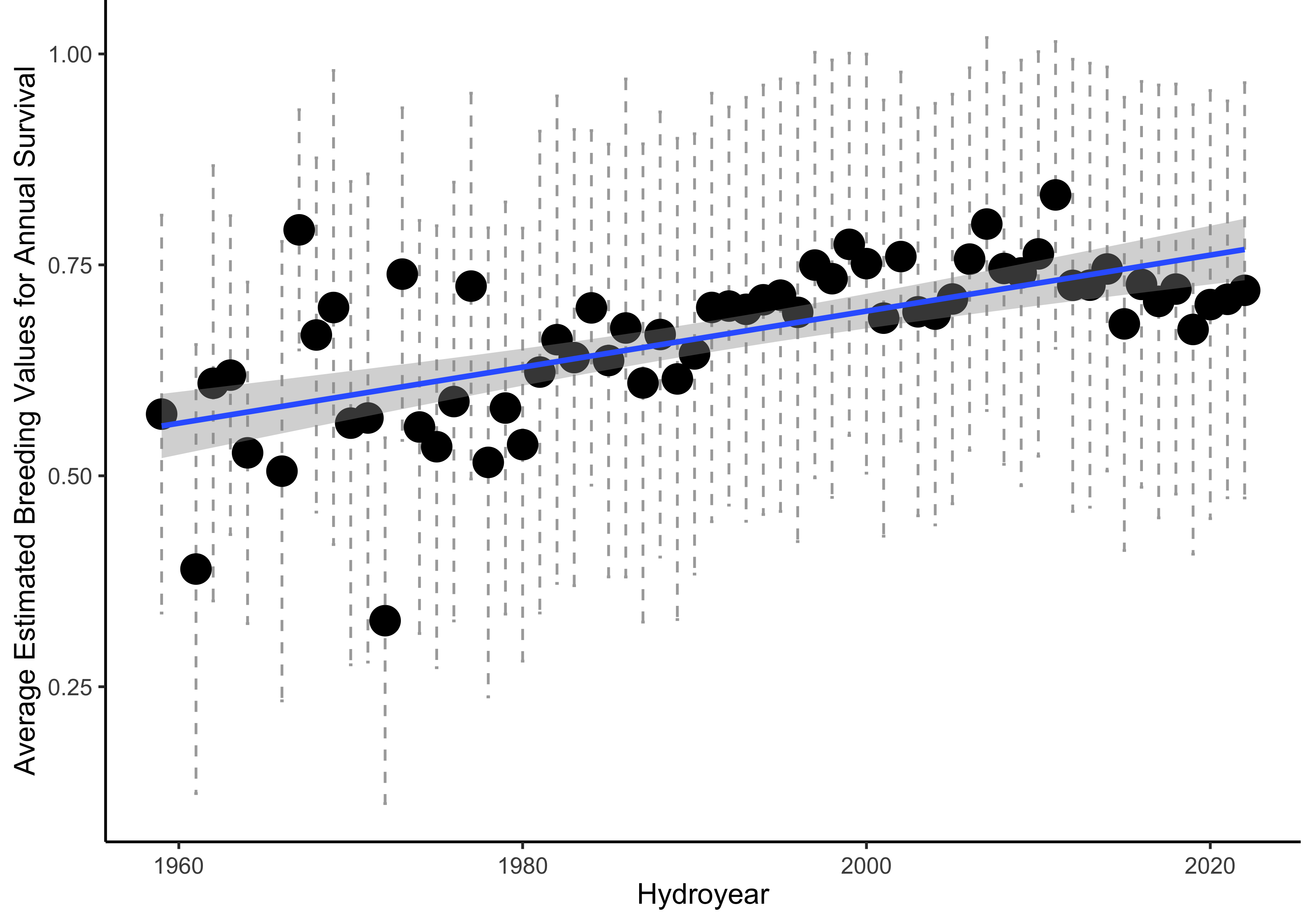

**Figure S4**. **Genetic trend for annual survival.** The estimated breeding values (EBVs) for annual survival are averaged within each cohort: we report the mean and standard deviation on the data scale as annual survival probabilities. Note that EBVs are only estimated as the “genetic merit” of the individual, i.e. the average phenotype that the individual would have if all phenotypic variance was genetic: for this reason, changes in EBVs do not represent actual phenotypic changes on the data scale.
